# Hippocampal representations of temporal structure increase in scale and symmetry across development

**DOI:** 10.64898/2026.03.10.710839

**Authors:** Owen W. Friend, Anthony M. Dutcher, Nicole L. Varga, Christine Coughlin, Alison R. Preston

## Abstract

Learning which experiences reliably co-occur in time is fundamental to episodic memory and improves markedly across childhood and adolescence. Although children and adults both engage the hippocampus while learning predictable sequences, the nature of the neural representations supporting statistical learning across development remains unknown. Here, we directly quantified item-level neural representations before and after children, early adolescents, and adults learned predictable temporal relationships between items, providing a direct measure of learning-related representational change. We identified three developmental shifts in hippocampal representation. First, although posterior hippocampus integrated temporally adjacent sequence elements similarly across age groups, integration of non-adjacent sequence elements increased with age in anterior hippocampus, indicating developmental expansion in the temporal scale of neural integration. Second, hippocampal representations changed in their directional organization, with children showing hippocampal representations reflecting only forward associations between adjacent events, whereas adolescents and adults exhibited bidirectional integration of sequence relationships. Third, functional connectivity between anterior hippocampus and frontoparietal cortex tracked statistical transition probabilities during learning and predicted memory performance. Together, these findings show that improvements in statistical learning during development reflect reorganization of hippocampal representations and hippocampal-cortical interactions, revealing how the developing brain constructs increasingly flexible representations of predictive temporal structure.

---

Statistical learning – the extraction of regularities from experience – allows individuals to detect which events reliably co-occur in time and to use those regularities to guide prediction (Saffran et al., 1996), supporting the ability to anticipate future outcomes and organize past experience. For example, a child repeatedly making a pizza for themselves may allow them to learn both directly adjacent relations (preheating oven must be done before cooking pizza) and longer-range dependencies (planning when to preheat the oven based on when the pizza should be ready). The hippocampus is thought to support this form of event linking by integrating temporally related experiences (Fortin et al., 2002), increasing the similarity of their neural representations to enable prediction (Schapiro et al., 2012; 2016). Indeed, hippocampal engagement during exposure to predictable sequences is evident even in infancy (Ellis et al., 2021). Yet behavioral evidence reveals protracted development of statistical learning; across middle childhood and adolescence, individuals increasingly link events separated in time and begin to flexibly generate predictions in both forward and backward directions (Arciuli & Simpson, 2011; Schlichting et al., 2017; Shufaniya & Arnon, 2018; Pudhiyidath et al., 2020; Forest et al., 2023a). If the hippocampus is engaged early in life, how do its representations of temporal regularities change to support continued gains in statistical learning? Here, we test the hypothesis that age-related improvements reflect increasing temporal scale and symmetry of hippocampal representations, as well as increasing neural sensitivity to transitions between events, as hippocampal subregions and their interactions with frontoparietal cortex mature into adolescence (Calabro et al., 2020; Langnes et al., 2020). Rather than examining these features in isolation, we directly quantify multiple component aspects of neural representation within the same paradigm, allowing us to test how they jointly contribute to developmental gains in statistical learning.

One fundamental feature of neural representation that may change across development is temporal scale – the window of time across which related elements are integrated in memory. Representational scaling has been a central organizing principle in animal models of hippocampal function and in adult neuroimaging work (Kjelstrup et al., 2008; MacDonald et al., 2011; Poppenk et al., 2013; Strange et al., 2014), yet never extended to the developing brain to understand how mechanisms of temporal memory emerge. Behavioral work suggests that adults link experiences across broader temporal windows than children (Pathman & Ghetti, 2014; Forest et al., 2023a). Specifically, even within experimental sequences unfolding over only a few seconds, children tend to remember events which appeared directly adjacent in time, whereas adults also link non-adjacent items (i.e., across intervening elements) that reliably co-occur. For example, after viewing shape sequences organized into consistent triplets (A-B-C), younger children (4–7 years) later recognized only directly adjacent elements (e.g., A-B), whereas older children (8-9 years) and adults remembered non-adjacent but temporally related elements that were never observed in sequence (e.g., A-C; Forest et al., 2023a). This pattern suggests that memory representations may expand in temporal scale with age, shifting from a reliance on direct observations to the integration of associations across time, even within short learning windows, let alone across hours or days (Forest et al., 2023b; Friend et al., 2026). Such differences would allow adults to predict consequences across broader temporal windows (e.g., rescheduling afternoon meetings after leaving late for work), while constraining children’s predictions to more immediate events (e.g., knowing one must take the next bus after missing the first), though this hypothesis has not been directly tested at the representational level.

Notably, adult neuroimaging work demonstrates that temporally related experiences are integrated in hippocampus, indexed by increased representational similarity (Schapiro et al., 2012, 2016; DuBrow & Davachi, 2014; Deuker et al., 2016; Bellmund et al., 2022; Pudhiyidath et al., 2022). Further, converging evidence suggests this temporal integration varies in scale along the long axis of the hippocampus (though see Deuker et al., 2016; Callaghan et al., 2021 for alternative findings). While anterior hippocampus can link events that span intervening items as well as broad temporal windows (even on the order of months; Nielson et al., 2015) to represent abstract, higher-order relations between experiences (Bellmund et al., 2022; Pudhiyidath et al., 2022), posterior hippocampus is thought to alternatively represent fine-grained, directly observed information (Poppenk et al., 2013; Brunec et al., 2018). Critically, while posterior hippocampus shows adult-like engagement by middle childhood, anterior hippocampus continues to mature into young adulthood (Ghetti & Bunge, 2012; Langnes et al., 2020), with emerging evidence that increasing functional maturity of anterior hippocampus may support integration across progressively broader scales of experience (Varga et al., 2026). Consistent with this protracted development, volumetric work has also linked anterior hippocampal maturity to statistical learning from ages 6-30 years (Schlichting et al., 2017). Because statistical learning requires linking information across time, the temporal window across which those links can be formed is a fundamental feature of the learning process. Here, we extend this work to direct tests of memory representation, predicting that children’s behavioral tendency to link only adjacent events in memory can be explained by the earlier-developing, discrete representational capacities of posterior hippocampus. Alternatively, the representation of non-adjacent (i.e., unobserved) associations in anterior hippocampus will emerge into young adulthood, corresponding to age-related behavioral gains in predictive memory.

While integration across broader timescales supports prediction across broader windows of time, a second mechanism we hypothesized would provide enhanced representational flexibility with age is symmetry - the linking of sequence representations in both the forward and backward direction in time (Schapiro et al., 2012). In adults, sequence representations are symmetrically integrated such that elements are linked in both forward and backward directions in time (Schapiro et al., 2012; Tarder-Stoll et al., 2024), and causal animal work suggests that hippocampus is required for backward associations specifically (Bunsey & Eichenbaum, 1996). This symmetrical linking provides flexibility by allowing one to remember either the next or the previous experience, depending on current environmental demands (e.g., when arriving late to your afternoon meeting, remembering that you left late that morning and planning your next morning differently). In line with this idea, the mature hippocampus selectively replays sequences in forward or backward order depending on context (Foster & Wilson, 2006; Diba & Buzsáki, 2007; Mattar & Daw, 2018; Wittkuhn et al., 2021), with forward replay linked to encoding direct observations and backward replay supporting the derivation of goal-relevant associations, even when backward associations are never directly observed (Davidson et al., 2009; Schuck and Niv, 2019; Gupta et al., 2010; Ambrose et al., 2016; Huang et al., 2018; Liu et al., 2021; Wimmer et al., 2023).

Importantly, behavioral work shows that children not only struggle to recall sequences in backward order relative to forward order, but are also less likely to derive temporal relationships that extend beyond their direct observations, with both abilities improving across development (Fivush & Mandler, 1985; Friedman, 1986; Friedman & Lyon, 2005; Jack et al., 2016; Ahmed et al., 2022; Friend et al., 2026). Furthermore, because the hippocampus maintains more stable representational states in adolescents and adults than in children, younger individuals may be particularly impaired at backward integration, which requires maintaining and comparing current event representations with prior states that are no longer directly present. In contrast, forward associations can be learned directly as sequences are experienced, placing fewer demands on representational stability over longer time intervals (Varga et al., 2026). Accordingly, we hypothesized that children would form asymmetric representations that encode sequences primarily in the forward direction in time (i.e., reflecting relations as they were directly experienced), whereas adolescents and adults would form symmetrical representations that additionally integrate sequences in the backward direction, representing temporal associations that were never directly observed.

Finally, beyond scale and symmetry which characterize how temporal relations are represented *within* sequences, we examined transition sensitivity – the brain’s ability to detect boundaries *between* sequences in continuous input – as a third mechanism underlying developmental gains in statistical learning. While structured input evokes hippocampal engagement in infants and adults at sequence boundaries (Ellis et al., 2021; Turk-Browne et al., 2009), behavioral work shows that the ability to acquire and deploy knowledge of transitional probabilities improves through adolescence and into adulthood (Pudhiyidath et al., 2020), suggesting additional neural mechanisms must support gains in transition sensitivity with age.

In adults, whereas hippocampus exhibits *both* sensitivity to boundaries and representation of temporal relations, cortical regions beyond the hippocampus are also implicated in predictive processing and the parsing of continuous experiences into discrete representations. Regions demonstrating such sensitivity to temporal boundaries include inferior frontal gyrus, lateral and medial prefrontal cortex, and lateral and medial parietal cortex (Schapiro et al., 2013; Hsieh & Ranganath, 2015; Cohn-Sheehy & Ranganath, 2017; Zhou & Turk-Browne, 2025). Adult work also demonstrates increased functional connectivity between prefrontal cortex and hippocampus during across-sequence transitions (DuBrow & Davachi, 2016). Accordingly, while hippocampus integrates experiences to represent underlying temporal structure, frontoparietal cortex may use this structure to segment continuous input and generate predictions about subsequent experiences (Hindy et al., 2016; Wang et al., 2017; Kok & Turk-Browne, 2018; Zhou & Turk-Browne, 2025).

Both hippocampus and frontoparietal regions mature through adolescence (Chang et al., 2016; Murty et al., 2016; Tang et al., 2020) and connectivity between anterior hippocampus and prefrontal cortex is among the latest to develop in early adulthood (Langnes et al., 2020; Calabro et al., 2020). Consequently, developmental differences in memory for temporal statistics may reflect age-dependent variation in interactions between hippocampus and cortex, particularly inferior frontal gyrus and medial prefrontal and parietal cortices, which have been demonstrated to represent temporal structure in adults (Schapiro et al., 2013; Pudhiyidath et al., 2022). We hypothesized that, with age, frontoparietal engagement and hippocampal-prefrontal coupling would increasingly track transitional probabilities between experiences, and that increased neural sensitivity to temporal structure would explain behavioral improvements in statistical learning across development.

Here, we provide an integrated account of how hippocampal representations of temporal structure reorganize across development by simultaneously quantifying their scale, symmetry, and sensitivity to transitions within the same task. Although prior work has linked hippocampal structure or activation at transition boundaries to developmental differences in memory, no study has directly measured how multiple component features of neural representation differ during childhood and early adolescence, a critical window for hippocampal maturation (Ghetti & Bunge, 2012; DeMaster et al., 2014). By indexing item-level representations immediately before and after learning, we isolate representational change itself, rather than relying on behavioral performance or univariate activation as proxies for the proposed representational mechanisms. This approach reveals how complementary hippocampal and frontoparietal mechanisms contribute to increasingly flexible statistical learning across development.

## Results

### Operationalizing temporal mechanisms

The present study examined how distinct representational processes jointly support temporal learning during a key developmental period in which both memory behavior and hippocampal and neocortical function undergo substantial change (7-12 years; Ghetti & Bunge, 2012; DeMaster et al., 2014; Calabro et al., 2020). Children (7-9 years) were compared to early adolescents (10-12 years) and adults (18-34 years) based on evidence that posterior hippocampus demonstrates adult-like activity around age 10, while anterior hippocampal function matures through adolescence (Ghetti & Bunge, 2012; DeMaster et al., 2014; Varga et al., 2025; 2026). Accordingly, we hypothesized that the scale and symmetry of neural representations would increase across this critical age range, and that developmental trajectories may differ by hippocampal subregion. Based on a power analysis (see STAR methods), thirty participants were recruited for each age group, with ten participants per year of age recruited for each developmental group. To jointly test multiple representational mechanisms across development, participants completed a statistical learning task while undergoing fMRI.

Participants first viewed all experimental stimuli in random order to allow for estimation of item-specific neural representations (**pre-exposure phase**; **Fig. 1A**). Arbitrary novel objects were used as stimuli to ensure neural representations were not confounded by pre-existing knowledge which may have varied between participants, obscuring developmental effects in the learning process. Next, during the key **triplet learning phase**, participants viewed the same items one at a time and were told only to “pay close attention and try to remember what you see.” Unknown to participants, items were organized with an underlying triplet structure such that four groups of three items each (triplets) were always viewed together in a fixed order (i.e., Item A, Item B, Item C; **Fig. 1.B.i**). Triplets were randomly arranged relative to each other, with the only constraint that the same triplet could not reappear until at least two other triplets had been displayed. As such, participants could learn that Item A, Item B, and Item C for each triplet were always presented in succession (i.e., A1→B1 and B1→ C1, transition probabilities = 100%). However, the transition from Item C1 to the next A item was randomized and less predictable (transition probability = 33.3%; **Fig. 1.B.i, 1.B.ii**). This triplet structure enabled representational analyses by allowing us to define adjacent and non-adjacent item pairs. Treating Item A as a reference, we directly compared integration for pairs of adjacent items (Item A to Item B) to pairs of non-adjacent items (Item A to Item C) which were never observed adjacent in sequence but shared the temporal context of the triplet (**Fig. 1.B.iii**). Finally, the unpredictable C to A structure across triplets allowed us to define boundaries between sequences for tests of transition sensitivity.

**Figure 1.**
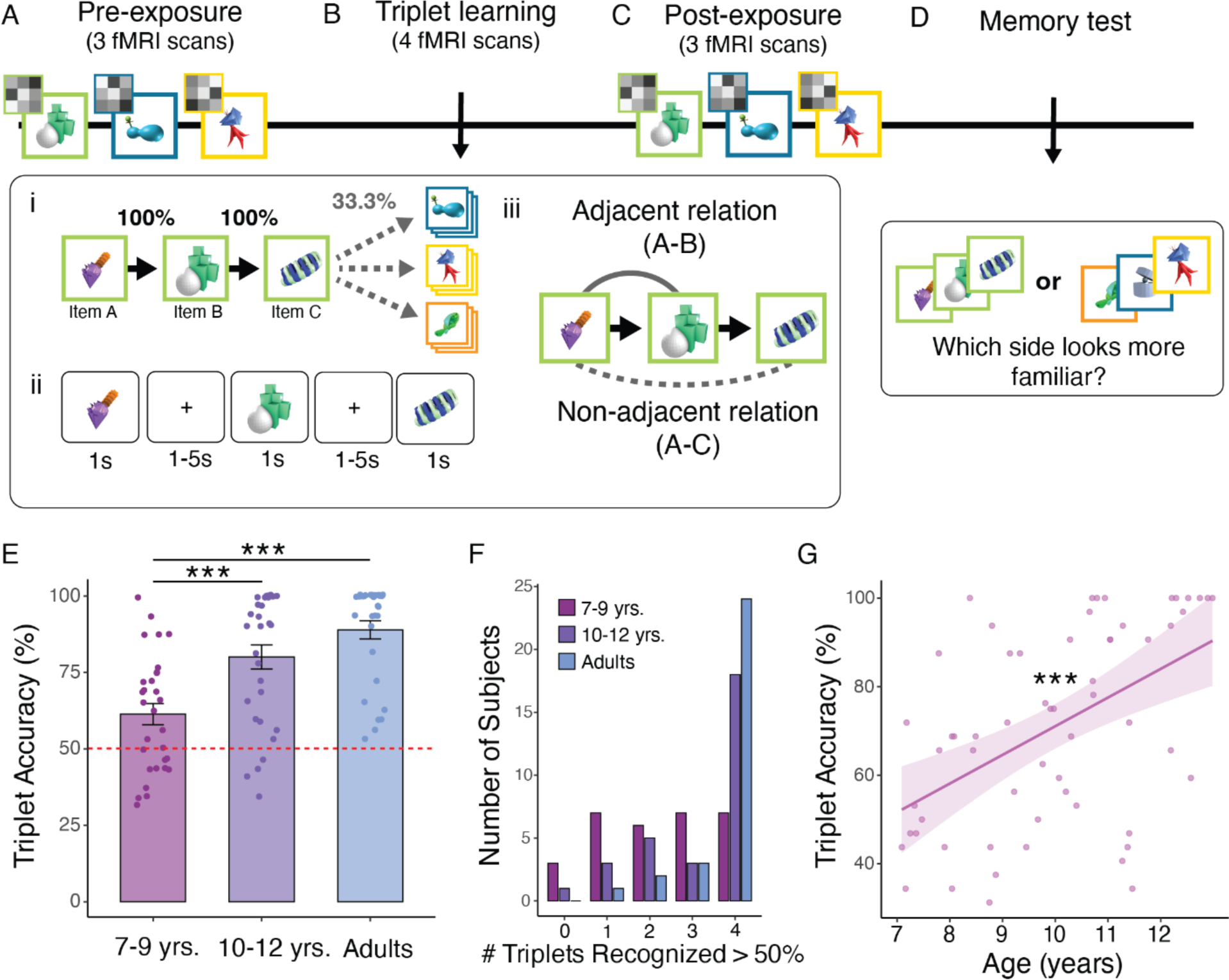
Statistical learning task overview and developmental gains in memory performance. Statistical learning task to assess representation of temporal regularities. **A**) During three pre-exposure runs, 12 experimental stimuli were presented in random order for 1s at a time. Stimuli were presented three times per run (nine times total). While viewing objects, participants engaged in a change detection cover task with high attention across age groups (see STAR methods). **B**) Following pre-exposure, participants engaged in triplet learning phase. **i**) Items were presented one at a time and participants were told to pay close attention and remember what they saw. Items were organized based on an underlying triplet structure such that groups of three items were always viewed in the same order. **ii)** Example timing of stimulus presentation during learning runs, with 1s object presentations followed by variable inter-stimulus intervals. Triplets were viewed four times per run, resulting in sixteen total presentations. Across-triplet order was randomly organized and could not reappear within two repetitions. **iii)** Triplet design allowed participants to associate adjacent items in sequence (observed adjacent relations (Item A-Item B) or non-adjacent items which were never viewed adjacent in sequence (Item A-Item C). **C**) Immediately after triplet learning, post-learning item representations were indexed identically to pre-exposure. **D**) Participants indicated memory for temporal triplets via two-alternative forced choice test, with preference for observed triplets suggesting statistical learning. Each triplet was paired with a foil item for 8 repetitions (32 total). **E**) Triplet-recognition accuracy by age group. All groups performed above chance (dashed line), with robust developmental improvements. **F**) Histogram of the number of triplets recognized above 50% accuracy for each age group. Younger children showed greater variability, but critically, nearly all subjects showed recognition of at least one triplet. Triplets were tested eight times each in random order, with different foil triplets as the alternative choice (see Methods). **G**) Continuous relation between age and statistical learning within developmental group. Linear trend indicates behavioral gains in statistical learning through childhood and early adolescence. Significance: *** *p <* 0.001.

After the key triplet learning phase (4 scanning runs, 24 repetitions of each triplet), participants completed three **post-exposure** scans (identical to the pre-exposure phase) to quantify representational change for adjacent and non-adjacent pairs of items as a function of seeing them in predictable order during learning (**Fig. 1.C**). Following post-exposure, participants completed a two-alternative forced choice test in which they repeatedly viewed one true triplet and one foil triplet and were asked which was more familiar (**Fig. 1.D.**, see STAR methods for additional detail). All foil triplets were composed of objects drawn from different triplets (e.g., A1, B2, C3). Thus, any recognition preference for the triplet over the foil provided evidence of knowledge of the underlying sequence structure.

### Behavioral evidence for sequence knowledge

Memory performance improved with age as adults (t(87) = 5.576, p < .001) and early adolescents (t(87) = 3.785, p < .001) each demonstrated higher recognition accuracy than children, while adults and adolescents did not significantly differ (t(87) = 1.791, p = 0.23; **Fig. 1.E)**, thus replicating prior work. Notably, all groups performed reliably above chance (child: t(29) = 3.272, p = 0.003; adolescent: t(29) = 7.619, p < .001; adult: t(29) = 13.054, p < .001; **Fig. 1**). All age groups also demonstrated equivalent attention across tasks (pre/post attention: F(2,86) = 1.745, p = 0.181; learning attention: F(2,83) = 2.194, p = 0.118)) which did not relate to accuracy (p = 0.33), suggesting the identified differences were specific to statistical learning and not sustained attention (see Methods, Supplemental **Fig. S1**). Moreover, though performance improved with age, almost all children (27/30) and adolescents (29/30) demonstrated recognition above 50% (chance) for at least one triplet (**Fig. 1.F**), suggesting nearly all participants learned at least some of the transitional probabilities. Finally, as expected, we identified a significant age-related increase in performance within the developmental group (B = 0.065, SE = 0.015, t = 4.439, p < .001) but no non-linear effects of age (ps > 0.296), suggesting a consistent and progressive improvement across the sampled age range (**Fig. 1.G**) and motivating our subsequent tests of the underlying neural representations.

### Neural representations are integrated across broader windows of time with age

The first representational mechanism we predicted would relate to developmental differences in temporal representation is the scale of time across which events are neurally integrated (i.e., represented more similarly in memory; **Fig. 2.A**). We hypothesized that earlier maturation of posterior hippocampus would support integration of directly observed adjacent relations across all ages, whereas integration of non-adjacent temporal relations in anterior hippocampus would emerge progressively across development (**Fig. 1B.iii**). Using searchlight representational similarity analysis approach within the hippocampus (**Fig. 2.A**), we identified a cluster in left posterior hippocampus in which adjacent items (Item A-Item B) were integrated consistently across age groups (**Fig. 2.B**; COG MNI coordinates (mm) *x, y, z* = −27, −33, −9, 34 voxels). We further validated the age-invariance of this effect by probing regions that showed integration across age-residualized subject-level maps, identifying the same cluster in left posterior hippocampus (see Methods). In contrast to age-invariant integration in posterior hippocampus, we identified a cluster in right anterior hippocampus in which integration of extended items (Item A-Item C) increased parametrically with age (**Fig. 2.C;** COG MNI coordinates (mm) *x, y, z* = 33, −14, −21, 37 voxels).

**Figure 2.**
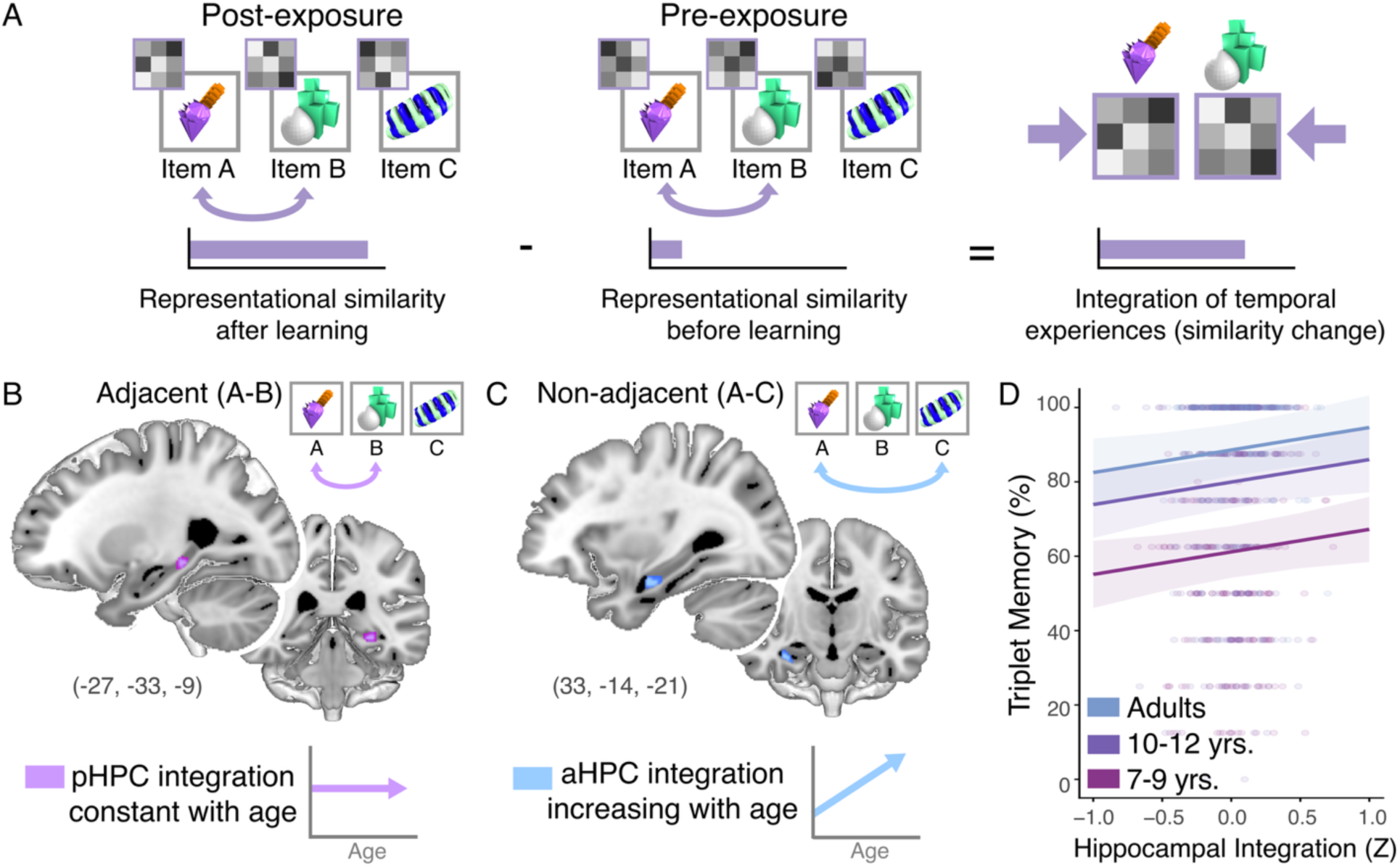
Anterior hippocampus integrates sequence elements across broader windows of time with age. **A)** Schematic of representational-similarity–based integration. For each pair of objects, we computed how much the voxelwise similarity between their neural patterns increased from *pre-exposure* (left) to *post-exposure* (right). Increased similarity from before to after learning reflects integration of temporal experiences. **B)** Hippocampus (light purple) demonstrated age-invariant integration of adjacent (A-B) items in sequence. **C**) With age, anterior hippocampus (light blue) demonstrates integration of extended (A-C) relations which require derivation of associations that are not directly observed, suggesting maturation of anterior hippocampal function supports memory for derived statistical relations. D) At the triplet level, the extent to which temporally-related items are integrated predicts behavioral memory performance. Each point represents one triplet from one participant (90 participants × 4 triplets = 360 observations), with integration predicting memory performance at the triplet level using a mixed-effects model. The horizontal bands of points reflect the discrete nature of the behavioral measure: because memory was assessed across eight trials per triplet, memory performance could only vary in increments of 12.5%. Regression lines illustrate the relationship between integration and memory performance for each comparison condition. This relationship was consistent across age groups and comparison types (adjacent vs. non-adjacent), suggesting that although development changes the temporal scale across which experiences are integrated, the relationship between integration and behavioral memory remains stable.

In addition to integration of non-adjacent experiences in anterior hippocampus emerging with age, we hypothesized that integration of temporally related items in hippocampus would correspond to memory above and beyond age, such that greater integration would support recognition of the learned triplets. In line with this prediction, we found that integration at the level of individual item pairs predicted memory for their triplets (B = 0.061, SE = 0.028, t = 2.145, p = 0.032). This effect was notably invariant of age or representational scale (adjacent vs. non-adjacent; ps > 0.09). That both adjacent and non-adjacent integration supported behavioral performance is consistent with the demands of the task, as memory for either adjacent or non-adjacent item pairs would be sufficient to distinguish a true triplet from a foil. Thus, in the primary model, hippocampal integration supported recognition similarly across adjacent and non-adjacent relations, while only neural integration of non-adjacent relations showed developmental change (see Supplement for exploratory analyses by hippocampal subfields).

Exploratory whole-brain searchlights additionally explored integration beyond the hippocampus (see Supplemental Results p. 1), identifying an age-related increase in A-B integration in medial prefrontal cortex (MNI coordinates x = 9, y = 60, z = −19; **Fig. S4**), with no additional age-related A-B or A-C effects, and no age-invariant A-B effects outside the hippocampus, consistent with recent developmental work localizing developmental change to hippocampus rather than MTL cortex more broadly (Varga et al., 2025). Accordingly, these findings suggest that hippocampal development increases the scale across which representations are linked, rather than changes in how integrated representations support memory behaviors.

### Hippocampal sequence representations shift in directionality from forward to backward with age

In addition to age-related increases in representational scale, we hypothesized that the directionality of neural integration would become more flexible with hippocampal development. Specifically, we predicted that children would integrate experiences only in the forward order (e.g., Item A cues Item B), consistent with representing their direct observations. We predicted that adolescents and adults, by contrast, would also link experiences in the backward order (e.g., Item B cueing Item A) even though they never viewed them in reverse, reflecting an enhanced ability to derive and integrate associations across time and form symmetrical temporal representations in hippocampus (Schapiro et al., 2012; Tarder-Stoll et al., 2024). To that end, we implemented a representational symmetry analysis from adult imaging work (Schapiro et al., 2012) which independently captures the extent to which a sequence element’s representation becomes more similar to the following element (forward integration) and to the preceding element (backward integration; **Fig. 3.A**).

**Figure 3.**
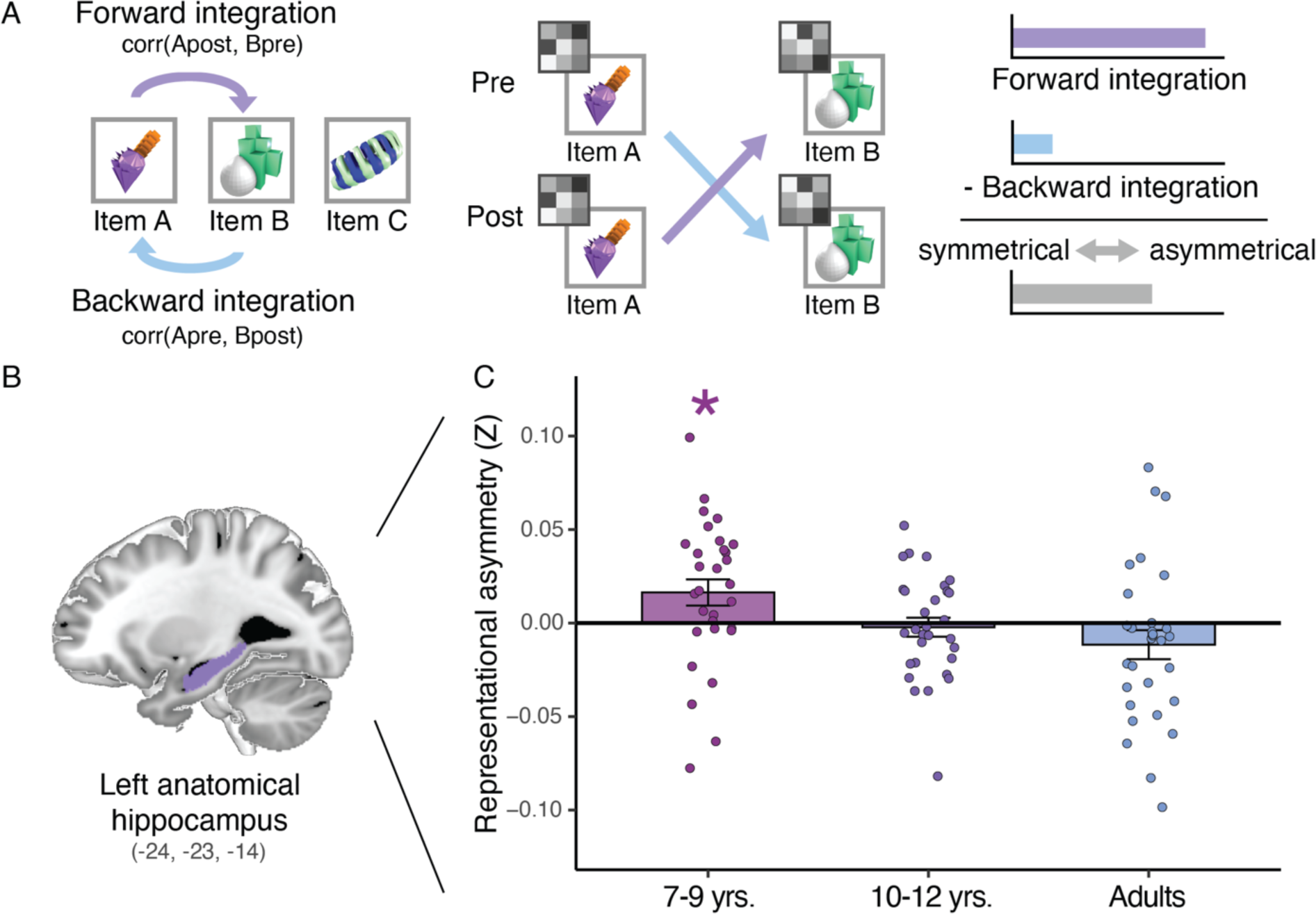
Symmetry of hippocampal sequence representations increases with age. Schematic of directional integration analyses **A)** Forward integration quantified how much the post-learning neural representation of an item (Item A) became more similar to the pre-learning pattern of the following item (Item B) (Schapiro et al., 2012). Backward integration measured the converse; similarity between the post-learning pattern of a later item (Item B) and the pre-learning pattern of the preceding item (Item A). Subtracting backward from forward integration indexes representational asymmetry, where higher values reflect stronger forward-biased integration, and values closer to zero reflect greater symmetry in how the hippocampus binds consecutively experienced items. **B**) Anatomical mask of the left hippocampus used for representational analyses (purple). **C**) Representational asymmetry (forward minus backward integration) across age groups. Younger children (7–9 years) showed significant asymmetric integration (a forward bias), whereas older children and adults did not significantly differ in forward and backward integration, suggesting bidirectional sequence representation. Significance: * *p* < 0.05.

Whereas we applied a standard searchlight approach to identify changes in representational scale, representational symmetry by definition reflects values approaching zero rather than discrete statistical thresholds because it captures the degree to which forward and backward associations are balanced (**Fig. 3A**; Schapiro et al., 2012). Moreover, because we identified representational changes in both anterior and posterior hippocampus and in both hemispheres from our representational scale analyses (**Fig. 2**), and because subregion differences in symmetry have not been reported in the adult or animal literature, we took the conservative approach of quantifying representational symmetry across the entire anatomical extent of the hippocampus. To avoid circularity (i.e., defining ROIs based on one representational metric and then testing a related metric in the same voxels), this analysis used anatomical regions rather than the searchlight clusters identified in the scale analyses.

In doing so, we identified an age group by hemisphere interaction (B = 0.028, SE = 0.013, t = 2.219, p = 0.027) that revealed developmental differences in representational symmetry for adjacent (A-B) item pairs in left hippocampus (the same hemisphere in which all groups demonstrated integration of adjacent items; **Fig. 3.B**). Within left hippocampus, children demonstrated asymmetrical coding, suggesting that they only integrated in the forward direction in time (A → B; t(29) = 2.327, p = 0.027). In adolescents and adults, forward and backward integration did not significantly differ, consistent with symmetrical representation (A → B and B → A; **Fig. 3.C**; adolescent: t(29) = −0.441, p = 0.663; adult: t(29) = −1.491, p = 0.147). Finally, because the current task only tested behavioral performance for sequences in the forward direction, we did not expect nor identify any links between symmetry and memory behavior (ps > 0.099; see Discussion).

### Hippocampal and frontoparietal cortical activity and connectivity track transition sensitivity with development

Finally, we predicted that developmental changes in transition sensitivity would reflect the joint maturation of hippocampus and its coordinated interactions with frontoparietal cortex, providing a mechanism that supports the representation of temporal relations not only *within* experiences, but also *across* them. Specifically, we predicted that 1) with age, neural activity in hippocampus and frontoparietal cortex would increasingly track transitions between experiences, and 2) with age, anterior hippocampus would increasingly coactivate with frontoparietal regions to prioritize sensitivity to the statistical structure of continuous experience. Whereas scale and symmetry measures reflect changes from pre-to post-learning, testing these hypotheses provided dynamic measures of how temporal relations are learned in real time.

To test how neural engagement varied with age during transitions between triplets, we implemented a univariate timeseries analysis with age as a parametric modulator to identify regions where activity at triplet boundaries increased with age, indicating greater sensitivity to the statistical structure of the input (**Fig. 4.A**). We identified several regions demonstrating increased transition sensitivity with age, including dorsolateral prefrontal cortex (dlPFC), inferior frontal gyrus (IFG), medial and lateral parietal cortex, posterior cingulate, and insula (**Fig. 4.B, Fig. 4.C**; Supplementary Table S1). To account for potential age differences in learning trajectories, we also compared neural engagement for early (Runs 1 and 2) and later (Runs 3 and 4) learning runs. In doing so, we identified a region in the hippocampal body where age differences in transition sensitivity were specific to early learning, potentially reflecting adults’ greater hippocampal sensitivity to transitions for new experiences. We did not find any regions that showed greater transition sensitivity during later learning.

**Figure 4.**
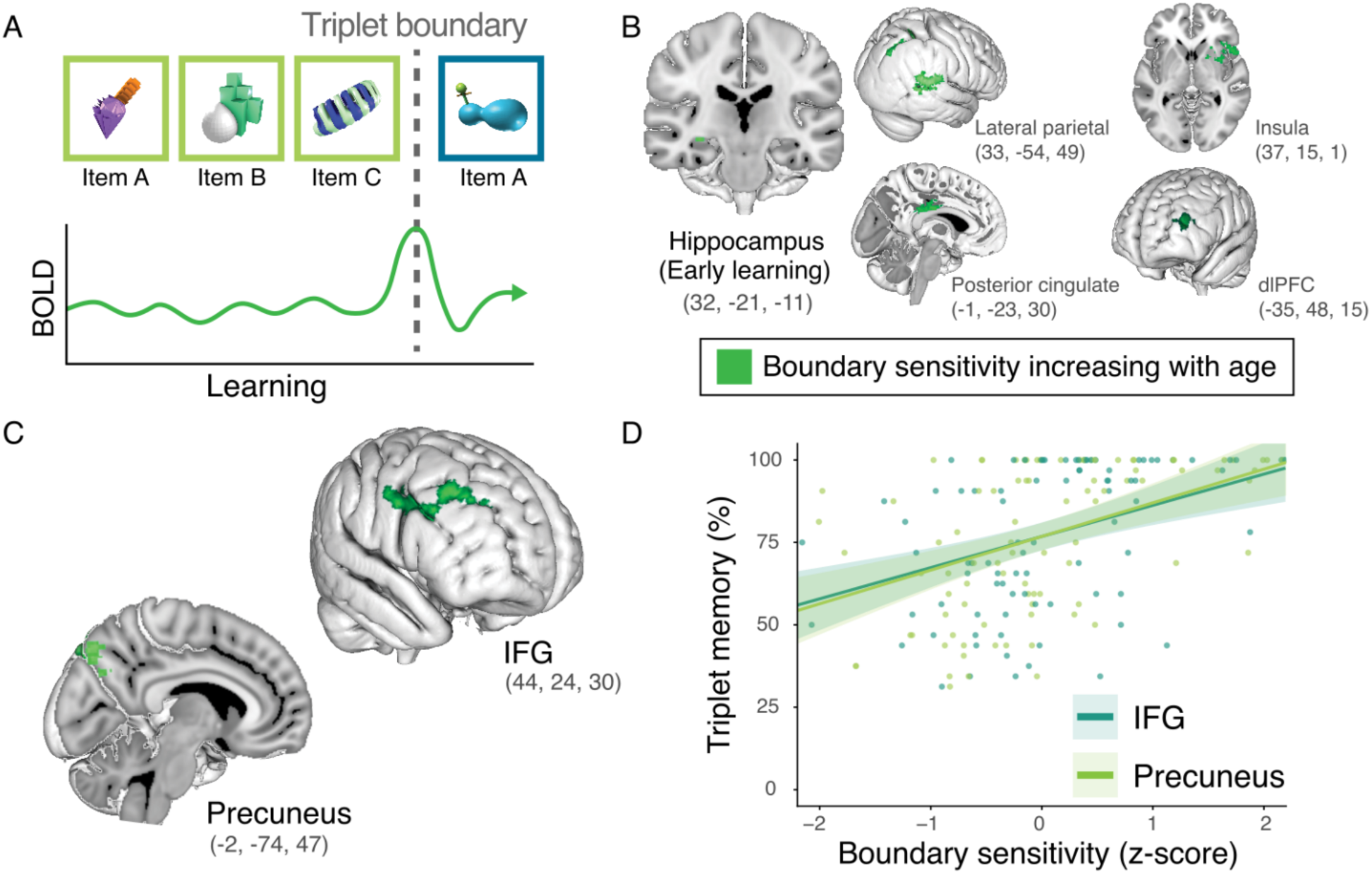
Hippocampal and cortical sensitivity to transitional probabilities support statistical learning. Univariate approach to quantifying age differences in neural engagement during triplet learning phase. **A**) First-level GLMs modeled transition sensitivity by identifying regions showing increased activation at triplet boundaries. Within the GLM, A Items were weighted as +1 while C items were weighted as −1, modeling increases or decreases in activation across the triplet boundary. B items were omitted due to the focus of this analysis on triplet boundaries. First-level GLMs additionally included motion and quality-related regressors. Second-level GLMs averaged activity across four learning runs, or compared Runs 1 and 2 (early learning) to Runs 3 and 4 (late learning). Third-level mixed-effects GLM from which clusters are derived was weighted by age (de-meaned) to reveal clusters demonstrating developmental change in engagement. **B**) Hippocampus demonstrates enhanced boundary sensitivity with age during early relative to late learning (Runs 3 & 4 > Runs 1 & 2). Lateral parietal cortex, posterior cingulate, and insula demonstrated enhanced boundary sensitivity with age across learning but did not track performance. **C**) Dorsolateral prefrontal cortex (dlPFC), inferior frontal gyrus (IFG), and precuneus demonstrated enhanced boundary sensitivity with age across learning. **D**) Above and beyond age, the extent to which IFG and precuneus showed enhanced engagement at triplet boundaries predicted performance on the task, revealing the behavioral correlate of cortical tracking of statistical structure. Points represent participant-level measures of boundary sensitivity in each identified region (IFG – blue; precuneus – green), plotted against the corresponding participant’s sequence recognition performance.

In addition to enhanced neural sensitivity to temporal structure, we predicted that this sensitivity would relate to behavioral differences in statistical learning. Consistent with this prediction, transition sensitivity in IFG and precuneus predicted memory performance above and beyond age (IFG: B = 0.0014, SE = 0.0006, t = 2.475, p = 0.016; precuneus: B = 0.0008, SE = 0.0003, t = 2.437, p = 0.017; **Fig. 4.D**), suggesting frontoparietal sensitivity to sequence transitions supports developmental gains in the ability to detect and remember predictive structure from continuous input.

Finally, we hypothesized that anterior hippocampal connectivity with cortex would become increasingly sensitive to temporal structure with age, based on prior work showing anterior hippocampal-prefrontal coupling at event boundaries in adults (DuBrow & Davachi, 2016) and evidence that anterior hippocampal projections to cortex are among the last to mature (Calabro et al., 2020). To that end, we implemented an age-weighted psychophysiological interaction analysis (PPI) using the anterior hippocampal cluster that showed increased integration with age (**Fig. 2D**) as the seed region (**Fig. 5A**). This analysis specifically identified regions in which older participants demonstrated enhanced functional connectivity with anterior hippocampus while viewing a triplet. We identified clusters in dorsolateral prefrontal cortex (**Fig. 5.B**; COG MNI coordinates (mm) *x, y, z =* −41, 37, −7) and inferior frontal gyrus (**Fig. 5.C;** COG MNI coordinates (mm) *x, y, z =* −47, 25, 22) which showed such enhanced anterior hippocampal coupling. Notably, no clusters showed enhanced coupling with posterior hippocampus, nor did anterior hippocampus show age-invariant coupling with any regions, suggesting this effect is specific to the maturation of anterior hippocampal connectivity. Additionally, no regions showed connectivity changes as a function of learning run, suggesting these effects were consistent across learning.

**Figure 5.**
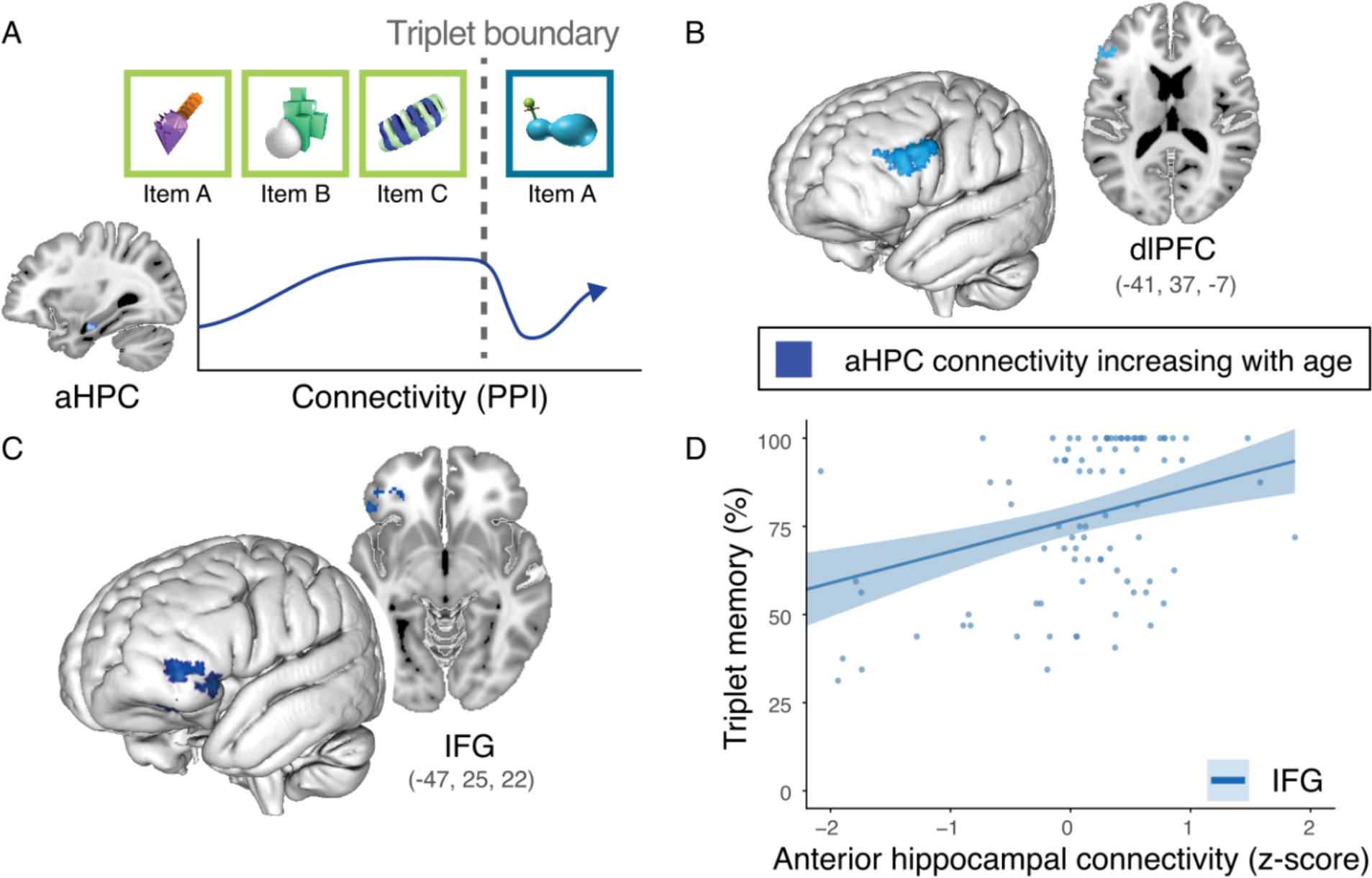
Emerging anterior hippocampal connectivity with prefrontal cortex supports statistical learning. Psychophysiological (PPI) approach to quantifying age differences in functional connectivity between anterior hippocampus and cortex during triplet learning phase. **A**) First-level GLMs modeled functional connectivity as interaction between eigenvariates of anterior hippocampal activity and item weights and durations. B and C Items were weighted as +0.5 while A items were weighted as −1 to capture connectivity for predictable transitions (A→B, B→C) over unpredictable transitions (C-A). Anterior hippocampal cluster revealing increased integration of extended pairs with age was used as seed region and back-projected into each participant’s native space before extracting eigenvariates. Second-level GLMs averaged across runs, while third-level mixed-effects GLM weighted connectivity maps by age (de-meaned) to reveal clusters demonstrating developmental change in functional connectivity. **B**) Lateral prefrontal cluster demonstrating increased coupling with anterior hippocampus during triplet presentation with age. **C**) Inferior frontal gyrus (IFG) cluster demonstrating increased coupling with anterior hippocampus during triplet presentation with age. **D**) Above and beyond age, the extent to which IFG showed enhanced functional connectivity with anterior hippocampus predicted performance on the task, suggesting anterior hippocampal connectivity with cortex coordinates both tracking of predictable transitions and subsequent memory for statistical structure. Points represent participant-level measures of aHPC-IFG connectivity at triplet boundaries, plotted against the corresponding participant’s sequence recognition performance.

Like transition sensitivity, functional connectivity between IFG and anterior hippocampus predicted memory performance above and beyond age (**Fig. 5.D**; B = 0.0011, SE = 0.0005, t = 2.098, p = 0.039), suggesting that anterior hippocampus not only represents extended relations between events (**Fig. 2.D**), but coordinates cortical activity to track and remember structured input. This result is consistent with our hypothesis that hippocampus both represents and tracks temporal regularities and, through coupling with frontoparietal regions, prioritizes frontoparietal sensitivity to this structure, together supporting prospection and memory for predictable experiences.

## Discussion

Statistical learning improves markedly across childhood and adolescence (Arciuli & Simpson, 2011; Schlichting et al., 2017; Shufaniya & Arnon, 2018; Pudhiyidath et al., 2020; Forest et al., 2023a), yet the neural mechanisms underlying these behavioral changes have remained poorly understood. The present study identifies three complementary representational mechanisms through which hippocampal and cortical systems support these developmental gains: changes in the temporal scale over which experiences are integrated, increasing symmetry in how sequence relations are represented, and enhanced sensitivity to transitions between sequences within hippocampal–frontoparietal networks. Although prior developmental work has primarily focused on general sensitivity to temporal transitions in infancy (Ellis et al., 2021), the present findings reveal age-related shifts in how temporal relations themselves are encoded. Specifically, whereas the earlier-developing posterior hippocampus represented temporally adjacent experiences similarly across age groups, the integration of non-adjacent experiences emerged across development in anterior hippocampus. Importantly, integration across both adjacent and non-adjacent timescales supported sequence recognition performance. In parallel, hippocampal representations became increasingly symmetrical across development, allowing sequences to be represented flexibly in both forward and backward directions. Finally, hippocampal engagement and connectivity with frontoparietal cortex became increasingly sensitive to transitions between sequences, with such transition sensitivity predicting improvements in statistical learning. Together, these findings provide a unified mechanistic account of how hippocampal representations and hippocampal–cortical interactions reorganize to support increasingly flexible learning from temporal structure across development.

We first examined how the temporal scale over which experiences are integrated in hippocampus changes across development. Animal models demonstrate systematic representational scaling along the hippocampal long axis, with anterior hippocampal representations spanning broader windows of experience (Kjelstrup et al., 2008; Poppenk et al., 2013; Strange et al., 2014). Consistent with this principle, adult neuroimaging studies show enhanced similarity of neural representations for temporally related experiences (Schapiro et al., 2012; Bellmund et al., 2022; Pudhiyidath et al., 2022) and evidence for differences in representational scale between hippocampal subregions (Collin et al., 2015; Nielson et al., 2015; Brunec et al., 2018). However, whether this fundamental scaling mechanism changes across development has remained unknown. Using representational similarity analysis, we found that posterior hippocampus integrates representations of temporally adjacent experiences similarly across children and adults, whereas integration across broader, non-adjacent temporal windows progressively increases with age in anterior hippocampus (**Fig. 2A–2C**). This developmental shift suggests that the mechanisms supporting broad temporal integration in anterior hippocampus are not fully expressed in childhood but gradually emerge across development. Notably, this temporal scaling parallels extensive spatial work across species demonstrating a representational gradient along the hippocampal long axis, with more precise local coding in posterior regions and broader, more abstract coding in anterior regions (Kjelstrup et al., 2008; Brunec et al., 2018). The presence of an analogous gradient for temporal relations suggests that representational scale may constitute a domain-general organizing principle of the hippocampus, emerging gradually across development rather than reflecting specialization for particular cognitive domains or mnemonic processes (e.g., space vs. time, encoding vs. retrieval).

Importantly, this gradient should not be interpreted as a strict functional dissociation, as anterior hippocampus can also represent detailed temporal and spatial relationships (Deuker et al., 2016), and developmental work has reported different trajectories of hippocampal computations depending on mnemonic demands (Callaghan et al., 2021). Differences across studies may therefore reflect the demands of the task and the information being represented in support of those demands. Our findings are best interpreted as a developmental shift in the relative temporal scale of hippocampal integration rather than strict functional division wherein detailed information is represented only posteriorly (Friend et al., 2026 for review). Whole-brain analyses further showed an age-related increase in adjacent pair integration in medial prefrontal cortex, but no additional age-related adjacent or non-adjacent pair integration effects and no evidence for age-invariant adjacent pair integration outside the hippocampus (**Fig. S4**). This pattern is consistent with recent developmental work localizing gains in associative and relational integration to the hippocampus rather than broadly across MTL cortex in children (Varga et al., 2026), while adult work suggests that associative representations can become increasingly distributed in cortex as structured knowledge is learned and reinforced (Kumaran et al., 2016).

Notably, integration at both temporal scales predicted sequence recognition (**Fig. 2D**), with no evidence that this relationship differed by representational scale. Because participants were required to recognize entire triplets, knowledge of either adjacent or non-adjacent pairings within a triplet was sufficient to distinguish true triplets from foils. Exploratory region-specific analyses further showed that posterior hippocampal integration of adjacent (A-B) relationships was associated with recognition accuracy above and beyond age, whereas anterior hippocampal integration of non-adjacent (A-C) relationships was not independently associated with memory accuracy (see **Fig. 2D**). Thus, the present task may be supported by adjacent associations without requiring the broader anterior hippocampal integration that emerges with age. This pattern suggests that development alters the temporal scale over which experiences are integrated, rather than whether hippocampal representations contribute to our simple measure of sequence recognition. The developmental emergence of broader anterior hippocampal integration may therefore explain age differences observed in tasks that more explicitly require integration across time. For example, memory for non-adjacent sequence relations improves into early adolescence (Forest et al., 2023a), and the ability to retrieve broader temporal context also shows protracted development relative to memory for adjacent order information (Pathman & Ghetti, 2014). Consistent with these findings, the present results suggest that children can rely on adjacent associations supported by posterior hippocampus, whereas broader integration across intervening experiences emerges with maturation of anterior hippocampal representations. While prior volumetric work proposed that developmental increases in hippocampal representational scale might drive improvements in statistical learning (Schlichting et al., 2017), the present findings provide direct representational evidence that integration across broader temporal windows emerges in anterior hippocampus through adolescence.

Critically, these findings do not imply that children are unable to acquire higher-level statistical regularities. Children can learn category-level statistics from item-level input by accumulating knowledge across repeated local relationships (Jung et al., 2021). Such learning may produce similar behavioral knowledge to memory linking across more extended temporal windows, but through a distinct mechanism. Specifically, children may build knowledge of broader relationships by repeatedly retrieving and linking locally learned associations across multiple experiences. For example, rather than directly representing the relationship between two items separated in time, a learner could retrieve a series of overlapping adjacent relationships and combine them to recover the broader structure. This iterative process can therefore support higher-level knowledge, but requires repeatedly accessing and combining intermediate associations on the fly rather than directly accessing a relationship that has been integrated across intervening experiences (Coughlin et al., 2026). Thus, developmental differences may reflect not only whether statistical structure is learned, but also how efficiently that structure can be organized and accessed across broader levels of abstraction.

Beyond changes in temporal scale, increasing flexibility of sequence knowledge may also depend on the emergence of symmetrical representations, whereby events are linked in both forward and backward directions in time. Causal animal work demonstrates that such symmetry is hippocampal-dependent (Bunsey & Eichenbaum, 1996). For example, rodents that learn that cue A predicts choice of B for reward also form the reciprocal association that B predicts A, supporting flexible behavior when task demands change. Following hippocampal lesions, however, animals fail to express these backward associations, resulting in reduced behavioral flexibility (Bunsey & Eichenbaum, 1996). Converging evidence from human neuroimaging similarly shows that hippocampal activity patterns reflect integration of sequences in both forward and backward directions (Schapiro et al., 2012; Tarder-Stoll et al., 2024). Consistent with this bidirectional integration, studies of hippocampal replay in animals and humans reveal both forward and backward replay following learning, with backward replay proposed to support the derivation of behaviorally relevant associations that were never directly experienced (Gupta et al., 2010; Ambrose et al., 2016; Huang et al., 2018; Liu et al., 2021; Wimmer et al., 2023). Because children often struggle to infer relationships beyond their direct observations, we predicted that the ability to integrate sequences in the backward direction, and thus form symmetrical memory representations, would emerge with age.

Applying this framework developmentally, we found that hippocampal representations after learning were asymmetric in childhood, reflecting only the forward associations that were directly experienced (A → B) (**Figs. 3A-3B**). By contrast, in early adolescence and adulthood we observed symmetrical coding, such that sequences were represented in both forward and backward directions (A → B, B → A) (**Fig. 3C**). This developmental shift aligns with theoretical accounts proposing that younger children’s memory is more constrained by direct experience, whereas associations between related events can be increasingly derived with age (Bauer, 2015; Shing et al., 2019; Friend et al., 2026). Accordingly, developmental increases in representational symmetry provide a plausible mechanistic account of behavioral findings showing that children struggle both to recall sequences in backward order (Fivush & Mandler, 1985; Friedman, 1986; Ahmed et al., 2022) and to infer temporal relationships that extend beyond directly observed events (Friedman & Lyon, 2005; Jack et al., 2016; Friend et al., 2026).

Notably, representational symmetry was not related to behavior in the present task. This pattern likely reflects the task demands, which required recognition of triplets in their experienced order. Because triplet memory was never tested in reverse, either forward or backward associations were sufficient to support recognition of a true triplet. Future work should therefore more directly test the behavioral consequences of representational symmetry using paradigms that independently isolate forward and backward integration across development, analogous to causal approaches in animal models (Bunsey & Eichenbaum, 1996). Interestingly, although neither older group showed evidence of asymmetry (**Fig. 3C**), we observed numerical decreases in forward coding alongside numerical increases in backward coding with age (**Supplemental Fig. S5**). We also found no association between representational symmetry and non-adjacent pair (A-C) integration in anterior hippocampus (**Supplemental Results p. 2**), suggesting that the developmental expansion of representational scale is not simply accompanied by greater bidirectional integration, but that these mechanisms are instead complementary, with both increasing with age. This pattern raises the possibility that representations following learning may increasingly prioritize derived associations over directly observed ones, a hypothesis that warrants targeted future studies.

Furthermore, developmental differences in symmetry were specific to adjacent item pairs (A–B pairs), despite the absence of age differences in the integration of these adjacent relations from pre-to post-learning (**Fig. 2B**). Thus, although adjacent experiences were integrated across all age groups, the directional structure of those representations changed across development. This dissociation indicates that representational scale and symmetry reflect distinct component mechanisms with separable developmental trajectories, emphasizing the importance of examining multiple dimensions of neural representation within the same paradigm. More broadly, these findings illustrate how new representational capacities can emerge within the same hippocampal circuitry across development, highlighting the value of extending mechanistic insights from animal and adult literatures to children and adolescents.

Beyond developmental differences in the scale and symmetry of within-sequence representations, a third complementary mechanism supporting statistical learning is sensitivity to transitions between sequences. Behavioral work suggests that detecting boundaries between experiences engages later-developing mechanisms than those linking elements within experiences, particularly during childhood and early adolescence (Coughlin et al., 2024). Thus, although neural sensitivity to temporal transitions is evident even in infancy (Ellis et al., 2021), the mechanisms that track these boundaries likely continue to mature with age. Consistent with this hypothesis, hippocampus and several frontoparietal regions showed greater increases in activity at across-triplet boundaries in older participants, indicating enhanced neural sensitivity to temporal structure with development (**Fig. 4**). Moreover, lateral prefrontal regions exhibited stronger functional connectivity with anterior hippocampus within triplets with age, suggesting the developmental emergence of coordinated hippocampal–cortical interactions that support tracking of temporal regularities (**Fig. 5**). Critically, differences in both transition sensitivity and hippocampal–prefrontal connectivity during learning predicted memory performance above and beyond age. Together, these findings suggest that although even infants detect temporal transitions, continued maturation of hippocampus, frontoparietal cortex, and their interactions support increasingly effective use of temporal structure to guide memory. The presence of early transition sensitivity therefore does not imply that other complementary mechanisms, including broader representational scale, symmetry, or hippocampal-cortical connectivity, are already mature.

While the sequences in the present task were relatively simple, they share key properties with event segmentation paradigms that demonstrate increasing neural sensitivity to structured, continuous input across development (Cohen et al., 2022; Benear et al., 2023). The present findings extend this literature by linking transition sensitivity directly to memory representations, revealing enhanced connectivity between prefrontal regions and the same anterior hippocampal area that exhibited developmental changes in temporal integration after learning (**Fig. 5**). Importantly, connectivity between anterior hippocampus and inferior frontal gyrus (IFG) predicted sequence recognition above and beyond age, underscoring the complementary roles of hippocampus and neocortex in representing temporal relations. One possibility is that cortical sensitivity to statistical structure tunes hippocampal representations to encode temporally structured experiences (e.g., sensitivity to triplet transitions guiding hippocampus to represent those triplets), with strengthening hippocampal–cortical interactions supporting increasingly structured memory across development (Hindy et al., 2016; Mack et al., 2016; Kok & Turk-Browne, 2018; Hauptman et al., 2024). By simultaneously quantifying neural activity, representational structure, connectivity, and behavior, the present study provides a mechanistic account of how hippocampal–cortical interactions support memory for continuous experience through multiple complementary mechanisms. Despite the use of the same anterior hippocampal region as the seed for the connectivity analysis and as the locus of developmental changes in non-adjacent (A-C) integration, connectivity did not covary with non-adjacent integration after accounting for age, suggesting that these measures capture unique sources of variance in predicting developmental differences in statistical learning.

Critically, the specific regions in which transition sensitivity and connectivity predicted memory beyond age—IFG and precuneus—were hypothesized *a priori* to support statistical learning based on prior evidence that these regions represent temporal structure in adults (Schapiro et al., 2013; Pudhiyidath et al., 2022). Moreover, these regions overlap with networks implicated in processing structured, continuous input across multiple cognitive domains, suggesting broader developmental mechanisms for tracking predictable structure (**Figs. 4C–4D, 5C–5D**). IFG development is often examined in the context of language, wherein structural maturation through adolescence has been linked to improved parsing of continuous input into meaningful units, such as segmenting speech into phonetic elements (Lu et al., 2007). By analogy, IFG may support temporal learning by facilitating the discrimination and grouping of sequential elements into structured units (i.e., triplets). In contrast, although the intrinsic anatomy of precuneus is relatively stable across development, its connectivity with default mode and frontoparietal networks continues to change through adolescence (Li et al., 2019), with such integration linked to memory retrieval processes in adults (Flanagin et al., 2023). Because precuneus also represents temporal structure after learning in adults (Pudhiyidath et al., 2022), its maturation may support the maintenance and updating of dynamic temporal representations as new relations are learned over time. Together, these findings suggest that maturation of IFG and precuneus supports the transformation of continuous input into structured, behaviorally meaningful memory representations.

Finally, the tightly controlled design of the present study enabled simultaneous tests of multiple representational mechanisms and facilitated comparisons across developmental and animal literatures. However, this control also imposes limitations. The use of novel objects was a strength because it reduced the possibility that developmental differences in prior knowledge or the ability to express that knowledge would be mistaken for differences in newly formed neural representations. At the same time, these stimuli and the short, deterministic triplet sequences are simplified relative to real-world experiences, which unfold over longer timescales and often contain probabilistic or hierarchical structure. Future work should therefore test how developmental changes in representational scale and symmetry generalize to more complex temporal environments and across familiar versus novel experiences (e.g., Schapiro et al., 2013; Pudhiyidath et al., 2020, 2022) as well as paradigms that more closely approximate real-world experience (e.g., Bellmund et al., 2022; Friend et al., 2026). Such comparisons will help determine how developmental changes in hippocampal integration mechanisms depend on the structure of experience.

In addition, although our behavioral measure was motivated by established work (Schlichting et al., 2017), tasks that separately assess forward and backward recall or require inference of missing elements from partial sequences could more directly test how developmental changes in representation shape memory behavior. Extending these approaches to continuous, multimodal paradigms such as movie viewing will further clarify how hippocampal functional maturation supports memory for complex experiences across development. Finally, because the study was cross-sectional, age differences in neural representation and performance cannot be entirely separated from cohort-related differences, underscoring the importance of longitudinal studies for directly tracking how representational scale, symmetry, and hippocampal–cortical interactions change within individuals across development.

Collectively, the present findings extend mechanistic insights from animal and adult literatures to human development, demonstrating that early hippocampal sensitivity to regularities gives way to fundamental shifts in how temporal relations are represented across childhood and adolescence. We identify coordinated developmental changes across three representational mechanisms, including the temporal scale and symmetry of hippocampal coding as well as increasing sensitivity to temporal structure within hippocampal–frontoparietal networks. Together, these findings show how developmental changes in neural representation and network interaction support increasingly flexible learning from temporal structure, providing a mechanistic account of how the developing brain extracts and organizes regularities from experience.

## Methods

### Participants

Participants were recruited such that 30 children (7-9 years), 30 early adolescents (10-12 years) and 30 adults (18-34 years) were included for representational similarity analyses. Within the developmental group, ten participants of each age were recruited. Age groups were approximately balanced by sex (child group: 21 female, adolescent group: 15 female, adult group: 15 female). Group sizes were selected based on a power analysis of the adult study from which our paradigm was adapted, which demonstrated hippocampal integration of items in sequence (d = 0.92 for two-sided t-test comparing hippocampal integration for within-sequence versus across-sequence item pairs; alpha = .05, 90% power; n = 11.63 per age group; Schapiro et al., 2012), supplemented by related developmental and adult work informing our choice of age ranges and broader recruitment target (Friedman & Lyon, 2005; Pathman & Ghetti, 2014; Ghetti & Bunge, 2012; Murty et al., 2016; Pathman et al., 2022; Schlichting et al., 2017; 2022). Age ranges for child and adolescent groups were selected as 7-9 years and 10-12 years based on developmental work suggesting that children ten and older tend to show adult-like engagement of posterior hippocampus during memory tasks (Ghetti & Bunge, 2012; DeMaster et al., 2014)

One hundred twenty-one total participants were recruited to achieve a final sample of 90 participants. Twelve participants were ineligible for scanning due to new dental work (2), new psychological diagnoses (2), English not being their first language (1), motion concerns during screening (1), aging out of the study’s age range (1), psychiatric assessment scores in clinical ranges (5) (see Behavioral screening session), while 18 withdrew following the behavioral screening session. In addition, one adolescent participant was scanned but fully excluded due to both technical difficulties and high motion during scanning, while one adult participant had one run of post-exposure excluded due to technical difficulties with the experimental task.

All participants were right-handed, native English speakers with normal or corrected-to-normal vision and hearing and without developmental diagnoses (e.g., Attention-Deficit/Hyperactivity Disorder, Autism Spectrum Disorder, etc.). Of the 60 developmental participants, 5% were Asian (3), 3.3% were Black/African-American (2), 71.7% were White (43), 1.7% were Pacific Islander (1), 16.7% were more than one race (10), and 1.7% declined to report race/ethnicity (1). Of the 30 adult participants, 23.3% were Asian (7), 6.7% were Black/African-American (2), 63.3% were White (19), and 3.3% were more than one race (1). 23.3% of developmental participants (14) and 36.7% of adult participants (11) identified as Hispanic. This community sample was socioeconomically and ethnically representative of the local Austin community. During both the behavioral screening session and MRI session, consent/assent were obtained using age-appropriate language in accordance with protocols approved by the institutional review board at the University of Texas at Austin. Upon completion, all participants received monetary compensation totaling $93 ($25/hour for 3.5 hours plus $5 bonus).

### Procedure

#### Behavioral screening session

All participants participated in a 1.5-hour screening session prior to scheduling an MRI visit. Following consent/assent, participants were provided with information about study logistics and MRI safety. Participants entered a mock MRI scanner to familiarize themselves with the environment and practiced holding still within the scanner. Next, participants (and their parents for minor participants) provided a series of measures to ensure eligibility for screening. Prior to scanning, all participants provided a series of measures to confirm eligibility. During screening, participants and/or their parents confirmed right-handedness, normal or corrected-to-normal vision and hearing, and native English speaking. In addition, general intelligence was assessed using the Wechsler Abbreviated Scale of Intelligence, Second Edition (WASI-II; Wechsler, 2011) vocabulary and matrix reasoning subtests. Participants scoring 2 or more standard deviations below the normed mean for their age were excluded. Psychiatric symptoms were assessed using the Symptom Checklist-90 Revised (SCL) (Derogatis & Cleary,1977) in adults and Child Behavior Checklist (CBCL) in children (Achenbach & Rescorla, 2001). Participants scoring in the clinical range (SCL: standardized global severity score > 62; CBCL: > 63) were also excluded from scanning. In addition to these assessments, participants and/or parents provided questionnaires related to socioeconomic status, pubertal stage, and life stressors as a part of separate studies. Finally, participants received instructions about the behavioral tasks they would perform during the MRI session and practiced condensed versions with alternate stimuli.

#### Experiment Overview

The experiment was organized into four primary phases. First, across three fMRI runs, participants viewed all experimental stimuli in random order to quantify baseline neural representations of each item (pre-exposure phase). Next, across four fMRI scans, participants completed the statistical learning task in which items were presented in temporally contiguous triplets (triplet learning phase) (Schlichting et al., 2017). Items within a triplet were always presented together and in the same order, while triplets themselves were randomly organized. After learning, participants completed three post-exposure fMRI runs wherein they viewed all stimuli in the same random order as during the pre-exposure phase to quantify learning-based representational change (post-exposure phase). Lastly, participants’ knowledge of the triplets was tested via a two-alternative forced choice triplet test outside of the scanner (triplet detection phase).

#### Materials

To ensure that neural representations of stimuli were not confounded by pre-existing knowledge or experience, we used 12 novel multi-colored 3D objects created in Blender (see **Fig. 1**; Schlichting et al., 2016). Serial positions and triplet membership for each stimulus were randomly assigned for each subject. Six additional novel objects were used during instructions and practice to familiarize participants to the task without exposing them to experimental stimuli. Stimulus presentation was implemented using custom MATLAB and Psychtoolbox-3 code (MathWorks, 2028; Kleiner et al., 2007).

#### MRI scanning session

To begin all MRI sessions, participants completed consent/assent and engaged in the same practice tasks as during the behavioral screening session. Immediately after entering the scanner prior to the pre-exposure phase, anatomical images were acquired while participants watched a self-selected movie.

##### Pre-exposure phase

To index learning-related representational change, we used a pre-post experimental design to quantify neural activity associated with each stimulus both before and after learning. Based on prior work suggesting that long fixed inter-trial-intervals result in strong item representation (Zeithamova et al., 2017), total trial durations were fixed at 4s. During each trial, an item was presented for 1s with a black fixation dot in the center of the image. After a random delay (50-275 ms) from the stimulus onset, the fixation dot changed to a black arrow facing left or right and remained on screen for 1s after stimulus offset. Participants were asked to quickly press one of two buttons on a button box based on which direction the arrow faced, providing a cover task to engage participants while quantifying item representations (Varga et al., 2025). Following the offset of the arrow, the black fixation dot was presented alone for 1.0s. The number and order of arrows facing left and right was randomized for each run.

Each run contained 36 trials (12 stimuli presented three times each) and 8 null trials (fixation dot only, no stimulus or arrow). Stimulus order was pseudo-randomized with the only constraint that items from the same triplet (see triplet learning) were not shown within two trials of each other. Participants completed three pre-exposure runs in succession, each lasting approximately 4 minutes, resulting in approximately 12 total minutes of pre-exposure scans including 9 presentations of each unique stimulus. Arrow detection performance was assessed as the proportion of trials for which the participant pressed the correct button to indicate the arrow’s direction.

##### Triplet learning phase

Across four fMRI runs, participants were repeatedly presented 12 items, organized into four temporally contiguous triplets (Schlichting et al., 2017). Items within a triplet were always presented in the same order (100% within-triplet transitional probability) while the triplets were presented in random order with the only constraint that the same triplet could not appear twice in succession (33.3% across-triplet transitional probability). By organizing items into triplets, we structured stimuli such that each triplet contained both adjacent temporal relations (Item A-Item B, Item B-Item C) and non-adjacent temporal relations (Item A-Item C). Critically, participants were provided no instructions related to recognizing the triplets. Rather, the only cue that the items were organized in triplets was the differences in transitional probability within and across triplets during learning.

All triplets were presented six total times during each run, resulting in 72 total learning trials per run. During each run of triplet learning, items were presented one at a time for 1s with a jittered inter-stimulus-interval (ISI) lasting 1, 3, or 5 seconds (29 trials, 29 trials, 14 trials respectively, randomly distributed). A black fixation cross was visible during the entire run and superimposed over each item during stimulus presentation. To ensure attention during triplet learning, participants were asked to indicate via button press when the fixation cross changed from black to red. Attention check trials lasted 1s each and appeared eight times during each run of triplet learning (32 total attention check trials).

To motivate and engage participants prior to triplet learning, we instructed all participants that they would be playing a game where they collected a stream of “alien jewels” across four planets. Each participant selected a character and spaceship and saw their character fly to one of four planets (corresponding to the four runs of triplet learning) prior to each scanning run. Participants were instructed prior to triplet learning and reminded between runs that their job was to pay close attention and remember the stream of jewels because they would be asked to remember the stream of jewels after “collecting” them by paying close attention. Because our primary question was whether participants could identify and remember statistical regularities from their experiences, participants were not given any information about the triplet structure or the structure of the triplet recognition test. Participants were also offered an effort-based five-dollar bonus for their attention during the triplet learning task.

##### Post-exposure item phase

To quantify neural representations of temporal relations between items, we assessed changes in memory representations elicited by triplet learning. To that end, participants completed a post-exposure phase identical to the pre-exposure phase so that representations could be compared between pre-learning and post-learning.

##### Triplet memory test

To index participants’ behavioral memory for triplets, participants completed a memory test outside of the scanner immediately following the post-exposure phase. The triplet memory test was organized as a two-alternative forced choice test in which participants were presented with a true triplet (i.e., seen during triplet learning) viewed in succession (Item A, Item B, Item C) and a foil triplet which was never observed during triplet learning and asked to indicate which was more familiar via button press. Foil triplets were comprised of items from three separate triplets which were presented in order by serial position (i.e. Triplet 1 Item A, Triplet 2 Item B, Triplet 3 Item C). True and foil triplets were randomly assigned to appear on the left or right of the screen and appeared one item at a time. The two triplet options were viewed sequentially rather than simultaneously, matching the item-by-item attentional demands of the learning phase. Items from the first option were presented sequentially 1s at a time. Then following a 1s delay, the three items from the second option were presented sequentially 1s at a time. Participants indicated via self-paced button press which option was more familiar (i.e., viewed during triplet learning). Participants could also repeat the triplet and foil presentation via button press as many times as desired to view the two options before responding. Participants rarely repeated trials, with average repetitions of 1.16 (SD = 0.15) in children, 1.12 (SD = 0.15) in adolescents, and 1.08 (SD = 0.09) in adults, with no age-group differences (ps > .218). During the Triplet memory test, participants completed 32 total choice trials (eight per triplet). Because all items were presented with equal frequency during both the triplet learning phase and the Triplet memory test, any familiarity preference for triplets compared to foils is indicative of memory for learned temporal relations.

### fMRI acquisition and preprocessing

Functional magnetic resonance imaging (fMRI) was collected while participants completed the pre- and post-exposure (6 runs) and triplet learning portions (4 runs) of the task.

#### Image Acquisition Parameters

Whole-brain imaging data was acquired on a 3.0T Siemens Skyra system (n = 22 subjects) and a 3.0T Siemens Prisma system (n = 68 subjects). Imaging protocols were designed to be as similar as possible on each scanner, and critically, temporal signal-to-noise ratio (tSNR) did not vary between scanners in the developmental group (B = 3.000, SE = 1.826, t = 1.641, p = 0.106) or the adult group (B = −0.566, SE = 2.027, t = −0.279, p = 0.782). Moreover, all reported models were tested with a categorical scanner covariate and with a continuous tSNR covariate, none of which affected any results, suggesting protocols across scanners were effectively balanced.

For all participants, a high-resolution T1-weighted MPRAGE anatomical image was collected for co-registration and parcellation. A high-resolution T2-weighted coronal image was acquired perpendicular to the hippocampus for improved parcellation and for subfield-level. Fieldmaps were acquired prior to each fMRI phase (pre-exposure, triplet learning, post-exposure) and applied to the following functional images to correct for magnetic field inhomogeneities. Functional data was collected with a T2*-weighted multiband accelerated EPI pulse sequence with a multiband acceleration factor of 3 and a GRAPPA factor of 2. EPI pulse sequences were designed to maximize temporal signal to noise ratio (tSNR) from the medial temporal lobe while maintaining voxel size precise enough to identify and segment hippocampal subfields. For full imaging parameters for all images acquired see Tables 1-2.

**Table 1.** 3T Prisma MR Image acquisition parameters. Image acquisition parameters for 3T Prisma (68 subjects collected).

| **Image** | **TR** | **TE** | **Flip angle** | **FOV** | **Voxel size** |
|---|---|---|---|---|---|
| T1w MPRAGE | 1.90s | 2.43ms | 9 deg. | 256 mm | 1.0x1.0x1.0mm |
| T2 Coronal | 13.15s | 84.00 ms | 150 deg. | 166 mm | 0.4x0.4x1.5mm |
| Fieldmap | 0.668s | 5.0/7.46ms | 5 deg. | 192 mm | 1.5x1.5x2.0mm |
| T2* Functional | 2.00s | 33.00ms | 77 deg. | 200mm | 1.7x1.7x1.7mm |

**Table 2.** 3T Skyra MR Image acquisition parameters. Image acquisition parameters for 3T Skyra (22 subjects collected).

| **Image** | **TR** | **TE** | **Flip angle** | **FOV** | **Voxel size** |
|---|---|---|---|---|---|
| T1w MPRAGE | 1.90s | 2.43ms | 9 deg. | 256 mm | 1.0x1.0x1.0mm |
| T2 Coronal | 13.15s | 82.00ms | 150 deg. | 166 mm | 0.4x0.4x1.5mm |
| Fieldmap | 0.668s | 5.0/7.46ms | 5 deg. | 192 mm | 1.5x1.5x2.0mm |
| T2* Functional | 2.00s | 30.00ms | 77 deg. | 220mm | 1.7x1.7x1.7mm |

#### Preprocessing of B0 inhomogeneity mappings

Three total fieldmap images were acquired, and for each fieldmap scan, a B0 nonuniformity map was estimated from the phase-drift map(s) measure with two consecutive GRE (gradient-recalled echo) acquisitions. The corresponding phase-map(s) were phase-unwrapped with prelude (FSL 6.0.4).

#### Anatomical image preprocessing

One T1-weighted (T1w) image was acquired for each participant. The T1-weighted (T1w) image was corrected for intensity non-uniformity (INU) with the ANTS N4BiasFieldCorrection function (Avants et al. 2008; Tustison et al. 2010). The T1w-reference was then skull-stripped with a Nipype implementation of the antsBrainExtraction.sh workflow (from ANTs), using OASIS30ANTs as target template. Brain tissue segmentation of cerebrospinal fluid (CSF), white-matter (WM) and gray-matter (GM) was performed on the brain-extracted T1w using fast (FSL (version 6.0.4), Zhang et al., 2001). Brain surfaces were reconstructed using recon-all (FreeSurfer 7.3.2; Dale et al., 1999), and a gray matter mask was estimated by combining and binarizing all cortical and subcortical parcellations, including hippocampus, for each participant (Morton et al., 2020).

In addition, one T2-weighted (T2w) image was acquired for each participant, aligned perpendicular to the hippocampus. Like the T1w image, the T2w image was corrected for intensity non-uniformity using N4BiasFieldCorrection (Tustison et al. 2010) and was isolated using BET brain extraction. The T2w image was used to segment hippocampal subfields via Automated Segmentation of Hippocampal Subfields with a custom atlas (Schlichting et al., 2019), for exploratory subfield analyses reported in the Supplement, and to refine the pial surface as a part of Freesurfer’s surface reconstruction (Dale et al., 1999).

#### Functional data preprocessing

Preprocessing was performed using fMRIPrep 23.1.3 (Esteban et al., 2019) which is based on Nipype 1.8.6 (K. Gorgolewski et al., 2011; 2018). For each of the 10 BOLD runs per subject, the following preprocessing was performed. First, a reference volume and its skull-stripped version were generated by aligning and averaging 1 single-band references (SBRefs). Head-motion parameters with respect to the BOLD reference (transformation matrices, and six corresponding rotation and translation parameters) are estimated before any spatiotemporal filtering using mcflirt (FSL, Jenkinson et al. 2002). The estimated fieldmap was then aligned with rigid-registration to the target EPI (echo-planar imaging) reference run. The field coefficients were mapped on to the reference EPI using the transform. BOLD runs were slice-time corrected to 0.955s (0.5 of slice acquisition range 0s-1.91s) using 3dTshift from AFNI (Cox & Hyde, 1997). The BOLD reference was then co-registered to the T1w reference using bbregister (FreeSurfer) which implements boundary-based registration (Greve & Fischl, 2009). Co-registration was configured with six degrees of freedom. First, a reference volume and its skull-stripped version were generated using a custom methodology of fMRIPrep.

Several confounding time-series were calculated based on the preprocessed BOLD: framewise displacement (FD), DVARS and three region-wise global signals. FD was computed using two formulations following Power (absolute sum of relative motions, Power et al., 2014) and Jenkinson (relative root mean square displacement between affines, Jenkinson et al., 2002). FD and DVARS are calculated for each functional run, both using their implementations in Nipype (following the definitions by Power et al. 2014). FD and DVARS values for each volume were used to perform motion-related quality assurance to ensure that no effects of interest were masked by motion artifacts, particularly in younger children where motion artifacts are more common. To that end, we identified all high motion volumes, defined as functional volumes with FD > 0.5mm and standardized DVARS > 1.5. Any run in which 1/3 or more of volumes were high motion was excluded from any further analyses (Varga et al., 2025). One subject was excluded based on this criteria, while all other participants contributed all six runs of pre- and post-exposure and all four runs of learning.

Across the included participants, children had an average of 14.4 high-motion volumes per pre-post run (SD = 10.6), adolescents 6.21 (SD = 8.13), and adults 0.88 (SD = 1.29), corresponding to approximately 13.8%, 6.0%, and 0.8% of the 104 volumes per run, respectively. Tukey-adjusted pairwise comparisons indicated significant differences between all age groups (adolescent vs. adult, p = .025; adolescent vs. child, p < .001; adult vs. child, p < .001). All available trials were retained for included participants. Because motion was modeled in the item-level neural pattern estimation described below as is considered best practice in estimating event-specific neural representations (Mumford et al., 2012), we did not conduct additional post-hoc motion-specific analyses.

Finally, all functional runs were skull-stripped using the brain mask masked derived from FreeSurfer (see above) and smoothed with a 4mm kernel using FSL’s SUSAN function prior to analyses.

### fMRI Analyses

#### Regions of interest

We defined a bilateral hippocampal mask using a manual delineation of the 1 mm MNI152 template in combination with a cytoarchitectonic atlas (Öngür et al., 2003). To separate anterior and posterior hippocampus, we delineated subregions based on the uncal landmark based on related developmental work (Riggins et al., 2015; 2018): the anterior hippocampus included gray matter anterior to the uncus, while the posterior hippocampus included the body and posterior portions located after the uncus. Importantly, although we used the uncus to delineate anterior and posterior hippocampal regions in a robust and reproducible manner consistent with the existing literature, all searchlight analyses were conducted across the entire hippocampus. Thus, clusters were identified independently of subregion definition, which was applied only after cluster identification to classify significant clusters as anterior or posterior. These anatomical hippocampal masks were then reverse-normalized into each participant’s native functional space using custom participant-specific ANTs-derived transformations. Exploratory subfield analyses were not used to define the primary hypotheses and are reported separately in the Supplement to ensure comprehensive reporting based on the adult literature (Schapiro et al., 2012).

For whole-brain analyses, we additionally derived a probabilistic grey matter mask in template space. All subject-specific grey matter masks (see above) were transformed into MNI152 1.7 mm space (matching the resolution of the functional EPI images) using nearest-neighbor interpolation and combined across participants. Voxels in which at least 50% of subjects contributed functional data were retained and binarized to create a final probabilistic grey matter mask.

#### Representational similarity integration analyses

##### Quantifying item-level representation

To quantify neural representations of temporal relations between events, we implemented representational similarity analyses (RSA) (Kriegeskorte et al., 2008) using PyMVPA (Hanke et al., 2009), Nilearn (Abraham et al., 2014), and custom Python routines. First, we estimated item-specific neural patterns evoked during the pre-exposure and post-exposure phases using event-specific General Linear Models (GLM) for each object using the Least Squares – Single method (Mumford et al,. 2012). To do so, we created event regressors for each of the 12 unique items in each run of the pre-exposure and post-exposure scans (3 timepoints per item per run) and convolved them with the double-gamma HRF in FSL. To rigorously control for motion, we additionally included fourteen nuisance regressors including six motion parameters and their temporal derivatives, framewise displacement (FD), and DVARS. This process resulted in a single voxelwise beta image for each of the 12 items in each of the 6 pre- and post-exposure runs (72 total images per participant) in each participant’s native T1w anatomical space. These images were used for all representational similarity analyses and allowed us to directly compare the representation of individual items before and after triplet learning.

##### Indexing pre-post representational change

After computing voxelwise beta images for each item, we asked whether representational similarity between items varied based on adjacent vs. non-adjacent temporal relations, and based on pre-vs. post-triplet learning. To that end, we extracted representational similarity values separately for adjacent and non-adjacent item pairs within triplets. For each comparison, we created correlation vectors to which we added similarity values calculated as r(item 1, item 2) and transformed to Fisher’s Z. Critically, we did not compare related items within the same run due to the inherent temporal autocorrelation of the BOLD signal, but only compared across runs within the same experimental phase (e.g., run 1 item 1 to run 2 item 2; run 4 item 1 to run 5 item 2) (Mumford, et al., 2012). We then averaged the pre-exposure and post-exposure vectors, resulting in single representational similarity values for each condition (adjacent and non-adjacent pairs) within each triplet in each phase for each participant. Learning-based representational change was then calculated for each comparison as r(post) – r(pre), resulting in single RS values for each triplet.

##### Searchlight analyses

To identify where in the brain such pre-post representation change was evident, we conducted searchlight analyses to examine the extent to which temporal relations were integrated following learning. Specifically, we asked where in the brain we would see significantly greater similarity between adjacent (A-B) or non-adjacent (A-C) items within a triplet compared to unrelated items across a triplet (e.g., triplet 1 item A - triplet 2 item B). Searchlights were conducted in bilateral hippocampus as an a priori region of interest based on prior developmental and adult work showing hippocampal integration of items in sequence (Schapiro et al., 2012; 2016; Bellmund et al., 2022), as well as across the whole brain in subject-specific gray matter masks (see above). Similarity contrasts were defined as described above in Pre-Post Representational Change and were calculated iteratively across the brain/hippocampus in individual searchlight spheres (r = 3 voxels). Importantly, hippocampal searchlights were conducted across the full bilateral hippocampal mask, with anterior and posterior anatomical masks used only after cluster identification to classify significant clusters along the long axis.

As in the RSA approach described above, items were only compared to items sharing the same phase but not the same run. Within each sphere we implemented a random permutation approach to test the statistical significance of the observed similarity change. The observed value was compared to a null distribution of values calculated by randomly permuting the similarity values for all items regardless of triplet membership. This approach was repeated 1000 times per searchlight sphere and resulted in a z-score reflecting the difference between the observed within-triplet similarity value and the mean of the null distribution of across-triplet comparisons. All searchlights were conducted in the functional space of each individual participant.

Group-level analyses were conducted using FSL’s *randomise*, which performs nonparametric permutation testing on GLMs. Individual participant searchlight z-maps were first aligned to MNI space using ANTs and then entered into a voxelwise GLM in *randomise*. For analyses relating age to representation, the design matrix included an intercept and a continuous age regressor (years and months; demeaned) to model linear age-related changes in pre-post representational similarity. Group-level similarity values were compared to a null distribution of 5,000 permuted samples to determine significance. For each of the 5,000 permutations, a random subset of participants’ z-maps had their signs reversed, and the full GLM (including the age regressor) was re-estimated, producing a null distribution for both the intercept and age effects. Significance was determined by comparing the observed voxelwise statistics to the permutation-derived null distribution, with threshold-free cluster enhancement applied to control for multiple comparisons. Positive age coefficients in the resulting maps thus indicated regions where pre-post representational similarity increased linearly with age. For analyses assessing age-invariant representation (e.g., **Fig. 2B**), the exact same *randomise* approach was implemented without the continuous age regressor, with the resulting statistics maps from this one-sample test indicating regions where pre-post representational similarity was significantly greater than the null across the entire sample. To supplement these analyses with a direct test of age-invariant effects, we additionally residualized participant age from each voxel’s RSA z-values across the full sample and entered the resulting residual z-maps into the same group-level one-sample analysis. This analysis therefore directly removed the linear contribution of age before computing group-level statistics, providing an explicit measure of the representational similarity change that was not attributable to age.

##### RSA Cluster/Small-Volume Correction

To determine cluster significance all group-level analyses, we performed cluster correction at the whole-brain level and small-volume correction within hippocampus. For both approaches, we conducted simulations to determine how frequently we would expect to identify clusters of specific voxel sizes based solely on chance. To do so, we first used AFNI’s 3dFWHMx method with the -acf flag to estimate spatial autocorrelation coefficients based on functional data in individual participant space. Specifically, we computed residual images to capture the variance in our data that was not associated with the events modeled in our GLMs (see above) and extracted autocorrelation coefficients from these residual images. Next, using those autocorrelation coefficients, we used AFNI’s 3dClustSim (Cox, 1996) to quantify the minimum amount of contiguous voxels needed to identify a statistically significant cluster at a voxel threshold of p < .01 and a cluster threshold of p < .05 (two-tailed, second nearest neighbor). Based on this approach, minimum cluster size was determined to be 16 voxels in hippocampus and 205 voxels in whole-brain grey matter.

#### Quantifying hippocampal representational symmetry and directional integration

We implemented a representational symmetry analysis from adult neuroimaging work which assessed representation of temporal statistics (Schapiro et al., 2012). As in the prior adult work, representational symmetry was computed only in *a priori* anatomical masks (i.e., anatomical hippocampus) which were back-projected into each subject’s native functional space. We did not compute symmetry within the searchlight clusters themselves, as those clusters were defined on the basis of integration and would therefore not provide an independent test of directional integration.

To quantify integration in the forward direction in time, we computed correlations between item-level representations for initial items after learning (Item A-post) and the succeeding item before learning (Item B-pre). Thus, if representation of Item A qualitatively shifted towards the representation of Item B (positive correlation), participants integrated representations forward in time. Conversely, if the representation of Item B after learning (Item B-post) qualitatively shifted towards the representation of the *preceding* item (Item A-pre), this correlation would suggest integration in the backward direction. Importantly, forward integration reflects representation of direct observations (i.e., items are linked in the order they were observed), whereas backwards integration reflects representation of derived associations (i.e., items are linked in an order in which they were never observed). Subtracting backward integration from forward integration provides a measure of representational asymmetry, with positive values reflecting a bias towards forward integration. Asymmetry values were computed within anatomical masks of left and right hippocampus and averaged across the four possible triplets, resulting in a single asymmetry index for each subject within each region of interest (Schapiro et al., 2012). Symmetry was quantified for both adjacent A-B and non-adjacent A-C item pairs; developmental differences in symmetry were observed only for adjacent pairs. Laterality effects were assessed via linear mixed-effects regression including hemisphere, age group, and their interaction term in addition to a by-subject random intercept, with each subject contributing asymmetry values from left and right hippocampus.

#### Timeseries analyses

##### Univariate activity analyses

To quantify participants’ sensitivity to temporal regularities during triplet learning, we implemented univariate GLM analyses using FSL’s FEAT (fMRI Expert Analysis Toolbox) version 6.0.4. First, individual models were fit to each triplet learning run for each participant. Each model included two primary regressors specifying timepoints in which participants were viewing the first and third items in all triplets, convolved with the canonical double gamma hemodynamic response function (HRF). To specifically establish brain areas that showed sensitivity to triplet boundaries, we employed a first-level contrast weighting the first item in triplets as +1 and the third item in triplets as −1 (Sherrill et al., 2023). Thus, by subtracting activity estimates during the third triplet item from activity estimates during the following first triplet item, we quantified differences in neural signal specific to triplet boundaries. We additionally included regressors of no interest to account for remaining variance including timepoints in which participants viewed the second item in a triplet or solely viewed the fixation cross, as well as six motion parameters and their temporal derivatives, framewise displacement (FD), and DVARS (Power et al., 2013). High-pass temporal filtering was also applied to each regressor (128s).

After modeling sensitivity to triplets within each learning run, we resampled the resulting images into MNI template space using ANTS and computed an average statistics image with each run weighted equally. We additionally compared early to late learning to capture neural activity specific to initial learning rather than maintenance of triplets in memory (contrast: Runs 1 & 2 - Runs 3 & 4. To establish age-related differences in boundary sensitivity, we implemented mixed-effects regression (FLAME 1) in Feat with age (in years and months; de-meaned) as a parametric modulator. Resulting statistics maps revealed regions in which boundary sensititivity increased with age (see **Fig. 4**). For approach to identifying significant clusters, see Cluster/Small-Volume Correction. Finally, all significant clusters were back-projected into each participant’s native functional space, and average within-participant values for boundary sensitivity were computed for each participant from each identified region of interest using FSL’s *fslstats* on each participants’ native contrast-of-parameter-estimate (COPE) images (Schlichting et al., 2022). These participant-level boundary sensitivity measures were then compared to task performance via multiple regression, with each participant contributing one observation of boundary sensitivity per region of interest to predict recognition accuracy, controlling for age.

##### Functional connectivity analyses

To test the prediction that hippocampus would show greater coupling with frontoparietal cortex during triplet boundaries, particularly later in development, we implemented psychophysiological interaction (PPI) connectivity analyses. PPI analyses were set up similarly to univariate time-series analyses, with an additional regressor to model hippocampal activity. Specifically, eigenvariates (measures of the variance explained by each principal component of the seed-region time series) for hippocampal activity were extracted for each run using *fslmeants* and included in first-level univariate models in interaction with event regressors. Event regressors indicated timepoints during which predictable transitions (i.e., Item A-Item B, Item B-Item C) occurred (weighted as +1) compared to timepoints when transitions could not be predicted (i.e., Item C-Item A; weighted as −1) to specifically assess functional connectivity while observing statistically predictable input. Second- and third-level PPI analyses were carried out identically to univariate analyses above. Within-participant functional connectivity values were compared to behavioral performance identically to the univariate analyses above.

##### Timeseries Cluster/Small-Volume Correction

Residual images for run-level models were separately extracted for each learning run from both univariate and connectivity analyses. Using these residual images, cluster/small-volume correction were performed identically to the representational similarity analysis searchlight results (see Representational similarity analyses). Based on this approach, minimum cluster size was determined to be 17 voxels in hippocampus and 188 voxels in whole-brain grey matter for univariate analyses, and 16 voxels in hippocampus and 189 in whole-brain grey matter for connectivity analyses.

## Notes

https://github.com/owenfriend24/temple

### Competing Interest Statement

The authors have declared no competing interest.

### Summary of Updates

Revised version has updated formatting of headings and no change to main text.

### Funding

- National Institutes of Health: R01MH100121
- U.S. National Science Foundation: DGE2137420

