## Supplemental Materials for "Hippocampal representations of temporal structure increase in scale and symmetry across development"

### Supplemental Results

#### Quantifying representation across both pairs of adjacent items.

Primary RSA results focused on a comparison between adjacent (A-B) pairs and non-adjacent (A-C) pairs, because we experimentally held the A item constant to directly contrast how it was integrated with items that reliably followed it in time (B vs. C), as a function of their temporal distance within the sequence. Although the B-C pair also constitutes an adjacent temporal relation, it does not share a common anchor item with the A-C pair that is predictably meaningful (i.e., only the C item is shared, and one can never predict what item follows a C item). B-C pairs, therefore, were excluded from primary analyses to preserve this controlled comparison of A-B versus A-C integration. For exhaustivity, however, we repeated the analyses across all adjacent pairs, including both A-B and B-C comparisons. Under this broader specification, no hippocampal or gray matter clusters survived cluster correction, indicating that the primary representational differences were specific to the temporal scale at which the initial triplet item (A) was integrated with predictably subsequent elements. Thus, including B-C pairs reduced overall integration effects but did not yield qualitatively different patterns, supporting our decision to focus on the theoretically motivated A-B versus A-C comparison.

#### Whole-brain representational scale analyses

To determine whether developmental differences in temporal integration extended beyond the hippocampus, we repeated the representational similarity analyses across the whole brain. Searchlight analyses were conducted using the same pre-to-post learning representational change approach as the primary hippocampal analyses. We tested for age-related differences in integration of adjacent A-B and non-adjacent A-C item pairs, as well as testing for age-invariant effects. Group-level age effects were assessed using the same nonparametric permutation framework described in the main Methods, with whole-brain cluster correction.

For adjacent A-B relationships, we identified an age-related increase in representational integration in medial prefrontal cortex (MNI coordinates x = 9, y = 60, z = -19; Fig. S4). No additional regions outside the hippocampus showed a significant age-related increase in A-B integration. In contrast, we did not identify any regions outside the hippocampus showing an age-related increase in integration of non-adjacent A-C relationships. We also did not identify any age-invariant A-B integration effects outside the hippocampus.

**Region-specific associations between integration and memory accuracy**

Although we did not identify interactions between age group and temporal scale (adjacent vs. non-adjacent) in our primary analyses, we conducted exploratory region-specific mixed-effects models to determine whether integration within the hippocampal subregions identified in the primary RSA analyses was independently related to triplet recognition performance. Specifically, these models predicted recognition accuracy from integration by age group in each subregion individually. In posterior hippocampus, integration of adjacent A-B relationships was significantly associated with recognition accuracy above and beyond age group, consistent with the primary effect illustrated in Fig. 2 ($B = 0.112$, $SE = 0.042$, $t = 2.655$, $p = .008$). In contrast, integration of non-adjacent A-C relationships in anterior hippocampus was not independently associated with recognition accuracy ($B = 0.011$, $SE = 0.047$, $t = 0.223$, $p = .824$).

Thus, posterior hippocampal integration of adjacent relationships may have supported recognition performance in the present task, whereas the broader anterior hippocampal integration that emerges with age does not provide an additional behavioral advantage on this measure. The absence of an independent relationship between anterior integration and recognition accuracy should be interpreted cautiously because either an adjacent or non-adjacent association within a triplet was sufficient to distinguish a true triplet from a foil. Thus, the task does not require explicit memory for non-adjacent relationships or inference across intervening events, and future work using tasks that more directly isolate these abilities will be important for determining the behavioral consequences of broader anterior hippocampal integration.

**Relationships between complementary hippocampal mechanisms**

The present study tested three complementary representational mechanisms which we predicted would relate to behavioral change across development. To understand how these mechanisms interact with each other, we tested whether the developmental mechanisms identified in the main analyses directly predicted one another. First, we examined whether greater representational symmetry was associated with greater integration of non-adjacent A-C relationships in anterior hippocampus. We did not observe a significant association ($B = -0.162$, $SE = 0.363$, $t = -0.446$, $p = .657$), suggesting that the broader representational scale that emerges with age in anterior hippocampus is not directly tracked by greater bidirectional integration. Moreover, developmental differences in symmetry were observed for adjacent pairs even though adjacent A-B integration itself did not differ by age. Together, these findings suggest that representational symmetry and representational scale reflect separable components of hippocampal temporal representation, contributing distinct sources of variance to developmental change.

We similarly tested whether anterior hippocampal-prefrontal functional connectivity during learning was associated with the magnitude of non-adjacent A-C integration in anterior hippocampus while accounting for age group. Neither anterior hippocampal-IFG connectivity ($B = 0.00020$, $SE = 0.00021$, $t = 0.968$, $p = .336$) nor anterior hippocampal-dlPFC connectivity ($B = 0.00059$, $SE = 0.00048$, $t = 1.225$, $p = .224$) was significantly related to non-adjacent integration. Thus, hippocampal representational integration and hippocampal-cortical connectivity appear to capture separable aspects of statistical learning rather than mapping onto one another in a simple one-to-one fashion.

#### Supplemental Figures

**Figure S1**

*Attention check trials do not vary by age group during pre-/post-exposure or learning*

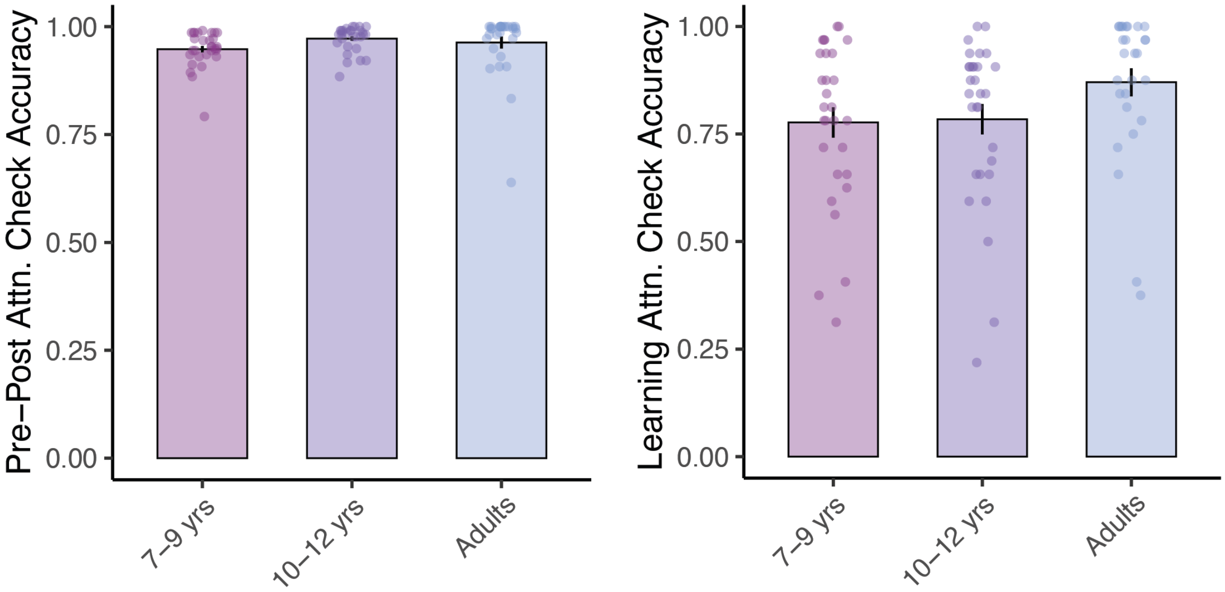


**Figure S1.** Age groups did not significantly differ in attention check accuracy during pre-/post-exposure ($F_{(2,86)}$ = 1.745, p = 0.181) or learning ($F_{(2,83)}$ = 2.194, p = 0.118). One adult subject excluded from pre/post attention analyses and three adults and one child excluded from learning attention analyses due to technical problems with MRI button box. Performance on attention checks was unrelated to triplet recognition (p = 0.33), further suggesting identified behavioral effects are specific to memory and statistical learning and not to attention.

**Figure S2**

*Pre-post integration of adjacent and extended item pairs in anatomical hippocampus and subregions*

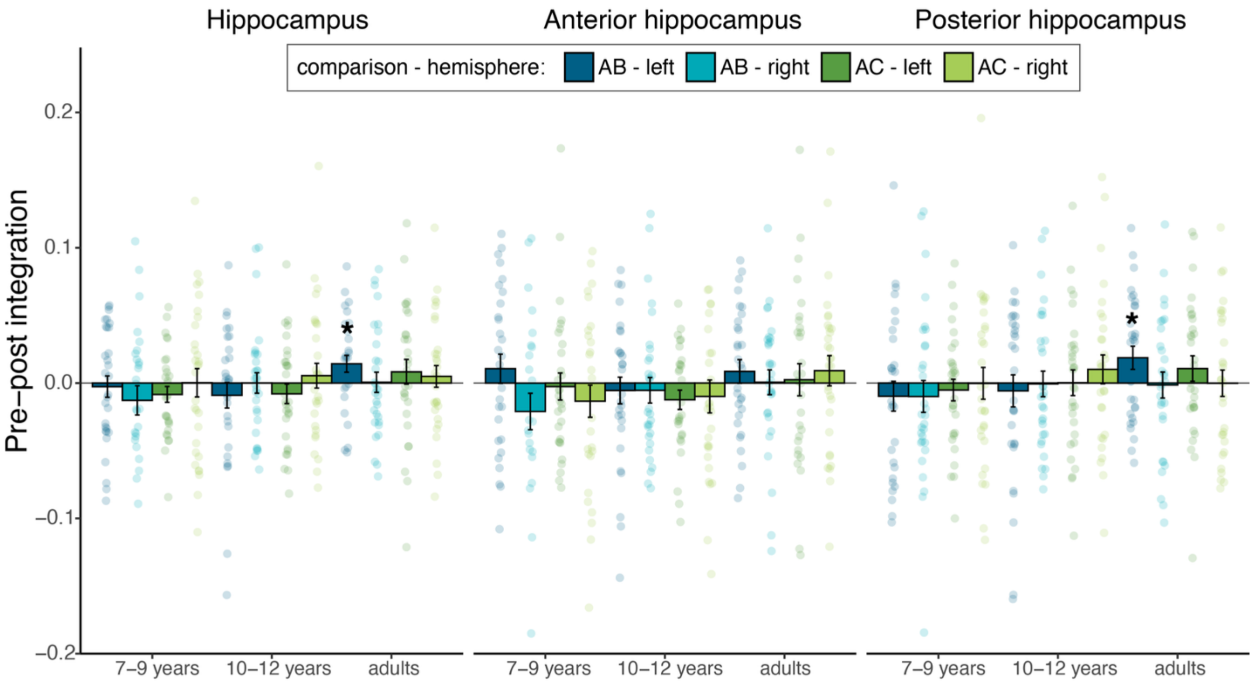


**Figure S2.** While primary analyses used standard searchlight approaches to identify clusters demonstrating neural integration, representational similarity measures within anatomical subregions are additionally reported for exhaustiveness. Mean pre-post integration by age group plotted for anatomical hippocampus (left panel), anterior hippocampus (middle panel), and posterior hippocampus (right panel). Bars indicate age group-level means for each comparison (AB, AC) and hemisphere (left, right), with error bars indicating SEM. Semi-transparent points reflect individual participant data (30 per age group for each comparison-hemisphere pair). Comparisons are color-coded (AB in blue, AC in green), with darker and lighter shades distinguishing hemispheres as indicated in the legend. Horizontal line at zero indicates no pre–post change, while asterisks denote integration statistically significantly greater than zero.

**Figure S3**

*Pre-post integration of adjacent and extended item pairs in anatomical hippocampal subfields*

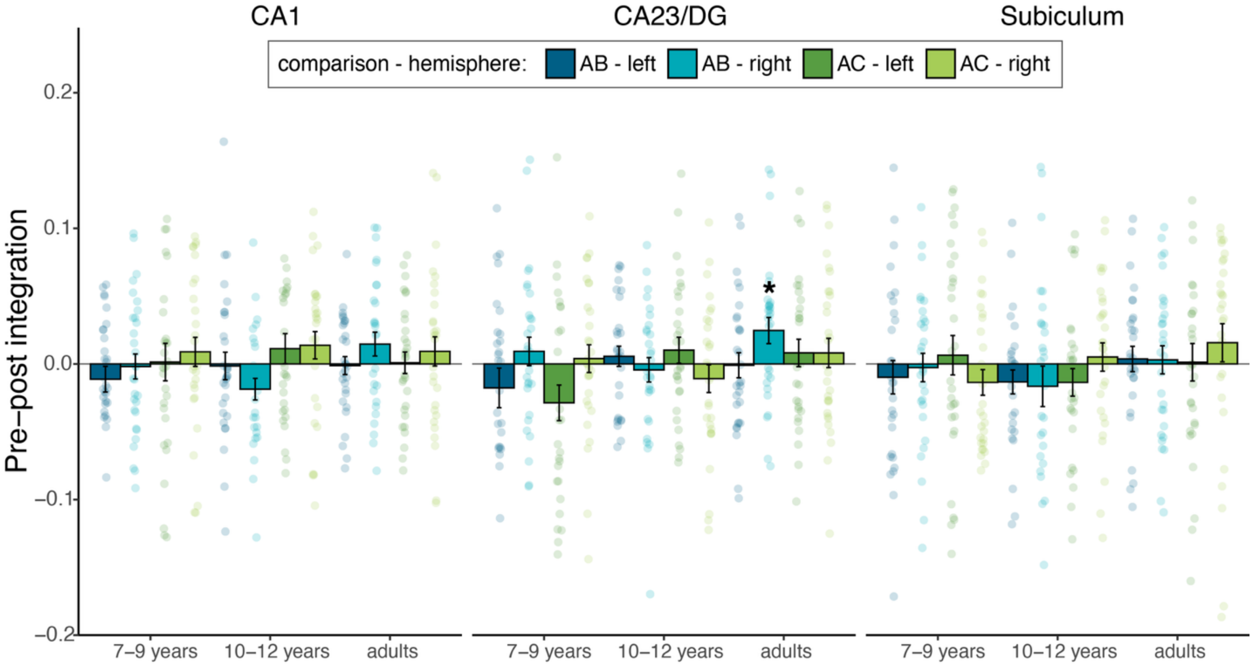


**Figure S3.** Primary analyses used standard searchlight approaches to identify clusters demonstrating neural integration, though here we additionally provide exploratory representational similarity measures within anatomically delineated hippocampal subfields for exhaustiveness. Mean pre-post integration by age group plotted for hippocampal subfield, including cornu ammonis 1 (CA1) (left panel), CA23/dentate gyrus (middle panel), and subiculum (right panel). Subfields are segmented individually for each subject from T1- and T2-weighted anatomical images (see STAR methods for imaging parameters) using ASHS automated segmentation (Yushkevich et al., 2014) with a previously-validated developmental atlas (Schlichting et al., 2019). Bars indicate age group-level means for each comparison (AB, AC) and hemisphere (left, right), with error bars indicating SEM. Semi-transparent points reflect individual participant data (30 per age group for each comparison-hemisphere pair). Comparisons are color-coded (AB in blue, AC in green), with darker and lighter shades distinguishing hemispheres as indicated in the legend. Horizontal line at zero indicates no pre–post change, while asterisks denote integration statistically significantly greater than zero.

**Figure S4**

*Medial prefrontal cortex increasingly represents temporal adjacencies with age*

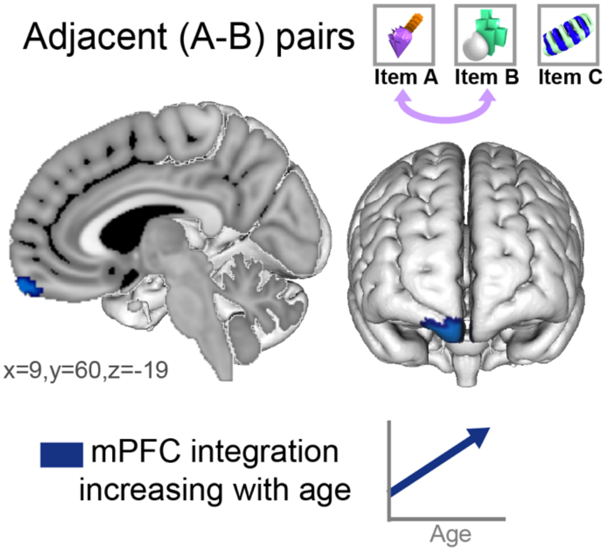


**Figure S4.** In addition to our primary hippocampal searchlight analyses, we conducted exploratory whole-brain searchlight analyses to identify cortical regions representing temporal structure. With age, a cluster in anterior ventromedial prefrontal cortex (dark blue) demonstrated increased integration of adjacent (A-B) relations. Thus, while hippocampus integrates adjacent temporal experiences across development (main text **Figure 2**), older participants increasingly also represent temporal adjacencies in medial prefrontal cortex. Unlike in hippocampus, prefrontal integration was unrelated to memory behavior in the present task.

**Figure S5**

*Hippocampal representations demonstrate numerical decreases in forward integration and increases in backward integration with age*

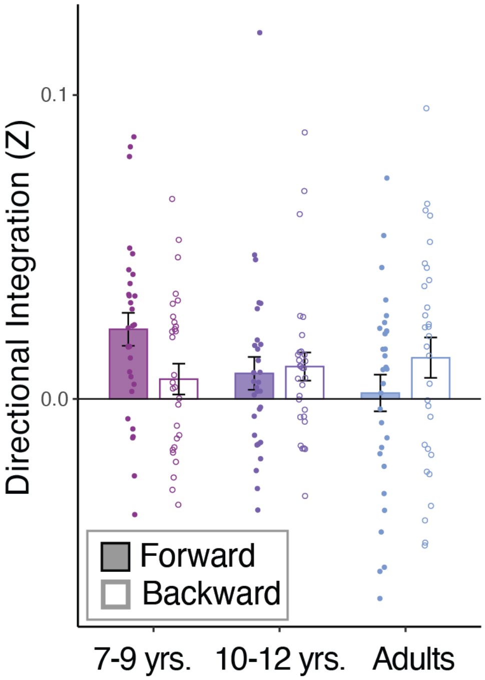


**Figure S5.** While children demonstrated evidence of representational asymmetry, forward and backward integration did not significantly differ in adults. For exhaustivity, we also compared integration in each direction to 0 using one-sample t-tests. Assessing each direction of integration separately, we confirmed as predicted that children integrated only in the forward direction in time (forward: $t(29) = 4.231$, $p < .001$; backward: $t(29) = 1.282$, $p = 0.11$). In early adolescents, by contrast, we identified significant or trending integration in both directions (forward: $t(29) = 1.561$, $p = 0.065$; backward: $t(29) = 2.307$, $p = 0.014$), suggesting integration of direct observations (i.e., A cuing B) as well as derived associations (i.e., B cuing A). Interestingly, adults only showed strong integration in the backward direction, (forward: $t(29) = 0.325$, $p = 0.37$; backward: $t(29) = 2.040$, $p = 0.025$; **Fig. 3.D**), possibly suggesting preferential integration for derived associations over direct experiences (see Discussion).

**Table S1**

*Regions demonstrating increased boundary sensitivity with age*

| | # Voxels | Hemisphere | X | Y | Z |
|---|---|---|---|---|---|
| Superior frontal gyrus | 1905 | bilateral | 2 | 4.7 | 61 |
| Postcentral gyrus | 1239 | left | -41 | -13 | 56 |
| Inferior frontal gyrus | 764 | right | 44 | 24 | 30 |
| Inferior frontal gyrus | 647 | left | -44 | 18 | 27 |
| Precuneus | 555 | bilateral | -2 | -74 | 47 |
| Angular gyrus | 447 | right | 61 | -45 | 20 |
| Insula/inferior frontal gyrus | 412 | right | 37 | 15 | 1 |
| Precentral gyrus | 309 | right | 44 | -13 | 61 |
| Lateral occipital cortex/angular gyrus | 282 | right | 33 | -54 | 49 |
| Dorsolateral prefrontal cotex | 223 | left | -35 | 48 | 15 |
| Posterior cingulate gyrus | 208 | bilateral | -1 | -23 | 30 |
| Insula/inferior frontal gyrus | 200 | left | -56 | 6 | 11 |
| Hippocampus (Runs 1&2 > Runs 3&4) | 17 | right | 32 | -21 | -11 |

**Table S1.** Cluster locations indicate center of gravity in 1mm MNI space
